# Riverine input and eddy edge effects on microeukaryotic biodiversity in the Northern Gulf

**DOI:** 10.64898/2026.07.20.739614

**Authors:** Sarah K. Hu, Alexis Adams, Abigail Day, Maggie Ellis, Norely Faz, Fernanda Haro, Madeleine Lerma, Kayla Nedd, Siddharth Seshampally, Meagan Sonsel, Chrissy Wiederwohl

## Abstract

Marine microorganisms drive the biogeochemical processes that sustain ocean ecosystems, such as primary production, nutrient cycling, and the transfer of carbon and energy to higher trophic levels. Single-celled eukaryotic organisms (microbial eukaryotes or protists) represent a multifaceted group that contribute to food web dynamics as primary producers, consumers, parasites, and nutrient remineralizers. The Northern Gulf of Mexico is a productive, river-influenced, semi-enclosed ecosystem with strong economic ties. To gain detailed insight into Gulf-based microbial communities, we present an 18S rRNA gene metabarcoding survey across 12 stations from the Louisiana coast to offshore Northern Gulf encompassing the surface to over 2,000 m. Together, water mass, the ratio of dissolved inorganic carbon to total alkalinity, distance to the coast, and depth structured protistan species composition; a secondary signal was associated with the edge of a Loop Current eddy. At the Mississippi River-Gulf interface, diatoms dominated the upper water column depths, with dinoflagellates, parasitic Syndiniales, and rhizaria making up the majority of the offshore communities throughout the entire water column. Shifts in species composition with Northern Gulf environmental gradients reflect varied trophic strategies and have implications for carbon transfer efficiency and food web structure. These results establish a baseline characterization of microeukaryotic biodiversity across coastal-to-offshore and surface to deep-sea gradients that provide critical context for future assessments of Northern Gulf ecosystem resilience.

## I. Introduction

Marine food webs represent a critical life-line for all living things in the ocean; this is particularly important for regions that are home to economies reliant on a healthy ocean ecosystem like the Northeastern Gulf bordering the southern coast of the United States (also known as the NE part of the Gulf of America or the Gulf of Mexico; hereafter referred to as “the Gulf”). Maintaining a healthy coastal ecosystem supports regional food security, protection from severe weather damage, regulation of climate conditions, and supports job security through commercial and subsistence fisheries and tourism (Lohrenz et al. 2008; Chakraborty and Lohrenz 2015). Marine microorganisms are the primary mediators of how biomass and nutrients enter, exit, and travel up the food web to larger organisms. In particular, single-celled microbial eukaryotes, which are also known as microeukaryotes or protists, play ecologically significant roles in marine food webs, acting as primary producers, consumers, and recyclers of organic material (Caron et al. 2012; Worden et al. 2015; Bachy et al. 2022). Protists are ideal ecosystem indicators; they are both biologically complex and respond rapidly to changes in their surroundings, making them representative of both animals (larger, multicellular eukaryotes) and microorganisms (Payne 2013; Berlinches de Gea et al. 2025). Thus, the study of environmental parameters that influence how microbial populations function in ocean ecosystems is essential for assessing and maintaining a healthy environment.

The Northern Gulf and Louisiana Shelf region is bordered by several sources of eutrophication, the largest being the Mississippi River. The fresh, nutrient rich riverine outflow from the Mississippi River is a dominant factor shaping the biogeochemistry of the entire Gulf and the subsequent microbial ecology, diversity, and distribution. Temperature, salinity, and nutrient gradients that form when the riverine input mixes with the NE Gulf water place strong selective pressures on both resident and introduced microbial communities. Mixed, high nutrient, regions close to the coastline typically favor fast growing species, like diatoms, while offshore sites harbor more diverse microbial communities (Wawrik and Paul 2004; Anglès et al. 2019).

Offshore, the Loop Current delivers warm water from the south through the Yucatan Straits (Sturges and Evans 1983). The confluence of weaker easterly winds in the Gulf, stronger easterly winds from the Caribbean, and surface warming, can result in an increased shedding of Loop Current eddies in the summer months (Chang and Oey 2012). Both warm- (anticyclonic) and cold- (cyclonic) core eddies separate from the Loop Current, subsequently enhancing or disrupting the stratified Northern Gulf region (Biggs 1992). Below the surface, the vertical structure of Gulf microbial communities is highly influenced by the oxycline, seasonal stratification, and the mixed layer depth (Rabalais et al. 2002; Wawrik et al. 2003). Eddy-induced upwelling and Ekman pumping can stimulate phytoplankton primary productivity at the surface, even in oligotrophic regions (McGillicuddy et al. 2007; McGillicuddy 2016). Microbial community species composition, trophic structure, and metabolic activity has been documented to be structured by mesoscale eddy dynamics (Harke et al. 2021; Gleich et al. 2024; Beatty et al. 2025).

Together with bacteria, archaea, and viruses, microeukaryotes mediate critical biogeochemical processes that sustain food webs and are well known to respond quickly to ecosystem changes. Prior Gulf-based analyses of microbial populations have focused on how bacterial communities shift seasonally or in response to storm or anthropogenic events (Dagg et al. 2007; Mason et al. 2016; Campbell et al. 2019; Henson and Thrash 2024), while non-HAB forming microeukaryotes and those below the surface have received comparatively less attention (Wawrik et al. 2003; Anglès et al. 2019; Campbell et al. 2019; Sidón-Ceseña et al. 2025).

A 2023 survey of microeukaryotic biodiversity and community structure spanning the Gulf coast to offshore and surface to deep sea, captured specific environmental niches microeukaryotes occupy. By pairing results from metabarcoding (18S rRNA gene sequencing) with concurrent physical and chemical parameters, we explore how environmental gradients structure protistan community composition across coastal-to-offshore and surface-to-deep transitions, how putative trophic and functional roles are partitioned across these gradients, and whether mesoscale features influence microbial community structure. We hypothesized that depth and proximity to the Mississippi River outflow are the primary factors organizing microeukaryotic biodiversity, with water mass and nutrient distribution as secondary factors. Water mass also emerged as a forcing factor alongside depth and coastal distance, indicating that these parameters co-vary rather than act independently. By resolving these relationships below the surface, this study establishes a depth-integrated baseline for the Northern Gulf.

## II. Materials & Methods

### A. Sample collection

Samples spanning three transects between the Southern U.S. coast (Louisiana, Mississippi River outflow) to offshore in the Northern Gulf (over 300 kilometers) were collected during the GoM GRADients 23 research cruise aboard the RV Point Sur, July 30 - August 4, 2023. The number of depths sampled at each station ranged between 2-12, where stations situated in deep-sea water had more samples collected (Table 1; Figure 1A). Seawater was collected via 10 L Niskin bottles mounted on a rosette above the CTD (SeaBird 19plusV2). Depths targeted for seawater sampling were chosen based on the real-time CTD downcast output data; typically Niskin bottles were fired at: 5-10 m above the seafloor to collect the deepest depth, 100-300 m above the seafloor, the depth of the subsurface oxygen minimum concentration (ml L^-1^), the subsurface salinity maximum, the deep chlorophyll maximum (DCM; subsurface increase in CTD fluorescence), and the surface (∼3 m depth). Discrete samples for molecular biology, microscopy, dissolved oxygen, dissolved inorganic carbon (DIC), total alkalinity (TA), inorganic nutrients, and salinity were obtained using methods outlined in the Go-SHIP Hydro Manual (Hood et al. 2010).

**Figure 1.**
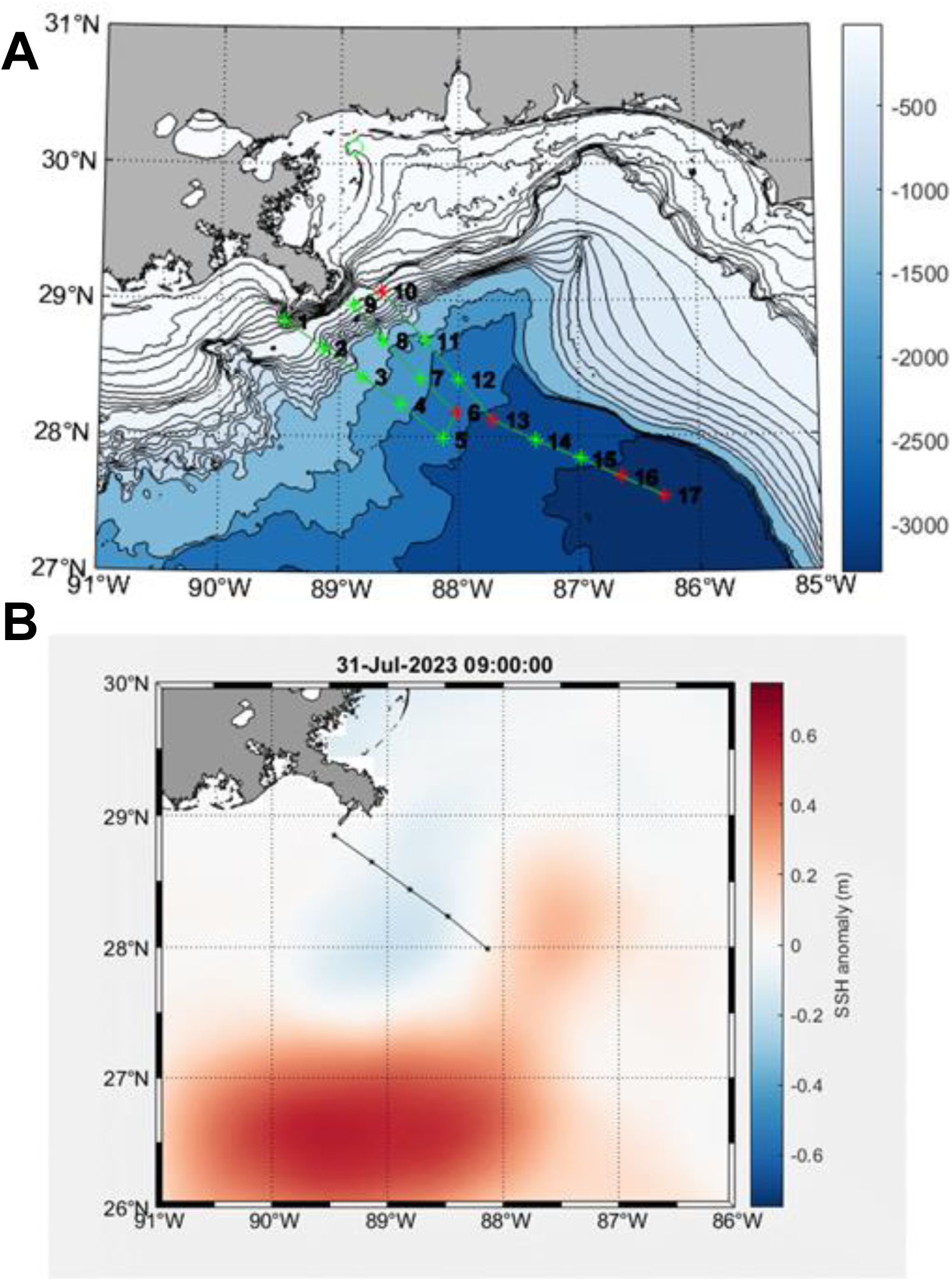
(A) Map of stations sampled during the Northern Gulf research expedition in July 2023. Color indicates depth in meters (m) and stations sampled in this study are in green. (B) Sea surface height anomaly (SSHa, m) during day 2 of the cruise survey (sampling of station 5). Animations for study area sea surface temperature and SSH anomaly can be found in supplementary materials.

**Table 1.**
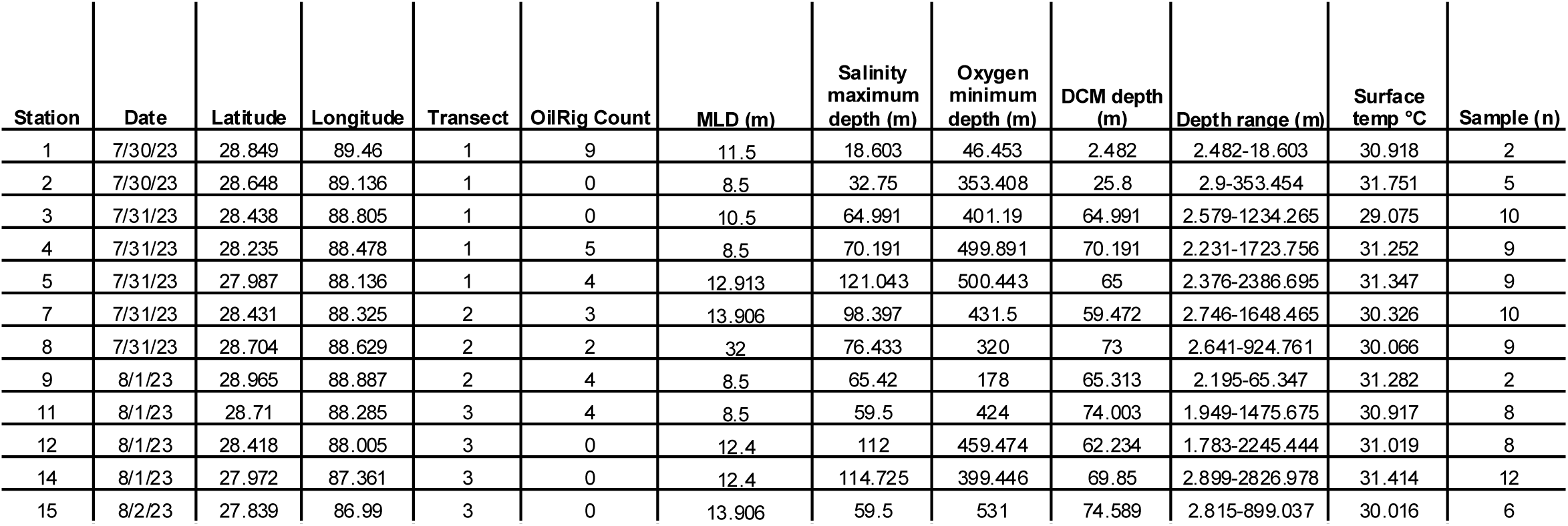
Station name, location, depth features, including Mixed Layer Depth (MLD) determined by Brunt-Vaisala, salinity maximum, and oxygen minimum, and deep chlorophyll maximum (DCM), range of depths sampled at each station, and the total number of depth layers sampled at each station.

Samples for molecular biology and microscopy were collected into acid-washed and cleaned 2-4 L bottles that were first rinsed with sample water 3 times and placed through a 200 µm woven mesh filter and kept in the dark (<45 minutes) until processing. A subset of sample fluid was preserved for eukaryotic and prokaryotic cell counts with glutaraldehyde (1% final concentration). Seawater for molecular analysis was vacuum filtered onto 0.2 µm PES filters; approximately 2 L was collected from euphotic zone samples and 4 L at subeuphotic zone depths (Table S1). Following filtration, filters were gently rolled into cryovials, RNAlater was added to each tube, and stored at −80°C.

Dissolved oxygen samples were collected directly from the Niskin spigot by overflowing the flask volume 3 times and then filling a glass flare-neck Erlenmeyer flask. Following collection, the sample was fixed with 1 mL of MnCl_2_ and then 1 mL of NaOH/NaI and capped with a ground glass stopper. Flasks were shaken vigorously to mix the solution. DI water was added to create a seal at the glass stopper. Samples were kept in the dark in the ship’s lab to acclimate to the lab temperature. Once acclimated, oxygen samples were analyzed using the Go-SHIP Winkler titration amperometric endpoint detection method. Thiosulfate was titrated to the flask to measure the endpoint. Dissolved oxygen was calculated using an amperometric technique (Langdon 2010).

DIC and TA samples were obtained by rinsing glass bottles 3 times, overflowing, poisoning each sample with mercuric chloride, and capping for room temperature storage until analysis. HDPE plastic bottles were triple rinsed and then filled with seawater. Seawater was filtered through a 0.2 µm PES membrane filter into 20mL HDPE plastic bottles and frozen at −20°C ahead of analysis. All nutrient samples were run by the Scripps Institute of Oceanographic Data Facility following Go-SHIP methods (Becker et al. 2020). Seawater was also collected from each niskin into borosilicate glass bottles double sealed with a thimble plastic insert and screw cap. Bottles were rinsed 3 times before filling to the neck and capping. Bottle salinity was measured using a Guideline 8400B Autosal with an accuracy of < 0.002 salinity. The Autosal was calibrated with IAPSO standard seawater daily.

### B. Regional assessment & physical oceanography

To characterize the Northern Gulf conditions, both sea surface height anomaly (SSHa) and satellite-based sea surface temperature (SST) were obtained from the National Oceanic and Atmospheric Administration’s Environmental Research Division’s Data Access Program (ERDDAP). SSHa and SST for every 3 hours between July 18-August 16, 2023 was extracted from HYCOM (HYbrid Coordinate Ocean Model) at 0.1 degree resolution (https://www.ncei.noaa.gov/erddap/griddap/HYCOM_reg1_latest2d.html; Figure S1, Supplementary animations).

Vertical profile features were determined using the CTD upcast data for the deep chlorophyll maximum (DCM), mixed layer depth (MLD), oxygen minimum layer, and the salinity maximum layer. All CTD data was processed using the Seabird SBE data processing software following the manufacturer recommendations for a SBE19plusV2 CTD system. CTD data was compared to both bottle salinity and bottle oxygen data and corrected using SBE application notes 31 (Application Note 31 Calculate Temperature and Conductivity Slope and Offset Correction Coefficients 2024) and 64.2 (Application Note 64-2 SBE 43 DO Sensor Calibration and Data Corrections 2024).

All large scale features were determined using the downcast information of the relevant CTD profiles, including DCM, MLD (temperature and salinity profiles), oxygen minimum (oxygen) and salinity maximum (salinity). Bottle depths were determined from the average depth of the center of the feature. Results from vertical profile features were visualized in Ocean Data View (ODV 5.8.5; Jan 27 2026). Water masses in the Gulf were determined following Portela et al. (2018), where temperature and salinity parameters associated with Antarctic Intermediate Water (AAIW), Caribbean Surface Water Remnant (CSWr), Gulf Common Water (GCW), North Atlantic Deep Water (NADW), North Atlantic Subtropical Underwater (NASUW), Tropical Atlantic Central Water (TACW), Tropical Atlantic Central Water nucleus (TACWn) were assigned to each sample. Samples that remained uncharacterized were separated by deep and upper water column uncharacterized water masses.

Mississippi (MS) River discharge data was obtained from the US Geological Survey National Water Information System data for the Mississippi River at Baton Rouge, Louisiana (U.S. Geological Survey, 2025), site: 07374000(U.S. Geological Survey, 2025). Sea surface temperatures (SST) from CTD data and station PSTL1 buoy, part of the NOAA National Data Buoy Center (1971), were used to classify the Marine Heat Wave index at time of sampling.

Oil platform data was acquired using the ArcGIS Online extension within ArcGISPro. This feature layer, updated on October 29, 2025, documents structures at or below the water surface designed for extracting and processing oil and natural gas. Data is curated and maintained by the United States Bureau of Safety and Environmental Enforcement, Office of Technical Data Management, Data Administration Unit. The number of oil rigs and relevant structures within 9 km of each station was documented as “oil rig count”. This distance captures both direct impact and the broader ecological footprint that may result from activities at the platform. A 9 km radius is large enough to capture downstream transport pathways and account for cumulative and compounding effects of contaminants while remaining at a spatial scale appropriate to distinguish platform-associated impacts from broader regional processes. Oil spill detection data was collected through NOAA’s Office of Satellite and Product Operations, which is managed by Environmental Response Management Application, NOAA Office of Response and Restoration, and the U.S. Environmental Protection Agency. Satellite images are used to estimate the area, level of confidence, and suspected point source of a potential oil spill.

### C. 18S rRNA gene metabarcoding

DNA was extracted from each sample using a modified lysis step ahead of the Qiagen DNA kit. Once filters in RNAlater were thawed on ice in the lab, the RNAlater was transferred to a separate tube and centrifuged for 5 minutes on high. RNAlater was decant and discarded; DNA lysis buffer was added to the tube to resuspend any leftover suspended genetic material. Along with additional lysis buffer, RNAse-free DNAse-free silica beads were added to a tube with the sample filter. Lysis buffer from the RNAlater tubes was transferred on top of filters and each tube was vortexed for up to 6 minutes. DNA products were quantified and cleaned using SPRI magnetic beads; the cleaned product was PCR amplified using primers specific to the V4 hypervariable region of the 18S rRNA gene: forward (5′-CCAGCASCYGCGGTAATTCC-3′) and reverse (5′-ACTTTCGTTCTTGATYRA-3′) primers (Stoeck et al. 2010). PCR reactions were conducted using 1X Q5 High Fidelity Master Mix (NEB #M0492S), 0.5 µM of the forward and reverse primers, and 1-2 ng of extracted, cleaned DNA. The thermal profile consisted of an initial step of 98°C for 2 minutes (min), followed by 10 cycles of 98°C for 10 seconds (sec), 53°C for 30 sec, 72°C for 30 sec, and 15 cycles of 98°C for 10 sec, 48°C for 30 sec, and 72°C for 30 sec, with an extension of 72°C for 2 min. Library preparation and sequencing was conducted using the Illumina platform NextSeq 2000 2×300 bp (University of Georgia Genomics and Bioinformatics Core).

### D. Data analysis

Amplicon sequences were processed using QIIME2 (Bolyen et al. 2019), where sequences were first quality controlled and primers removed with cutadapt (error rate = 0.1, overlap minimum = 3 bps; Martin (2011)). Using the DADA2 plugin available through QIIME2, paired end reads were truncated, used in a model to screen for errors (max-ee = 2), chimeras were removed (pooled method), and Amplicon Sequence Variants (ASVs) were determined (Callahan et al. 2016). Reference sequences for each ASV were assigned taxonomy using the PR2 database v.5.0.0 (Guillou et al. 2012; Vaulot 2021), using an 80% identity threshold in vsearch (Rognes et al. 2016). Laboratory blank samples were sequenced alongside study samples and used to reduce potential contamination using ‘decontam’, where ASVs with higher prevalence in blanks relative to study samples at a probability threshold of 0.5 (Davis et al. 2018). To further quality control the data, samples had to have at least 40,000 sequences and included ASVs were required to have ≥ 100 sequences across the entire dataset. For most analyses replicates were averaged ahead of normalization and data analyses. Taxonomic assignments from PR2 were filtered to isolate “Eukaryota” at the Domain level, removing ASVs labeled as “Bacteria”, “Unassigned”, or “Eukaryota:nucl”. ASVs assigned to the Opisthokonta taxonomic group were also removed, as they were considered outside the scope of this study and require alternate databases for appropriate taxonomy assignment. Within the PR2 taxonomy assignment, names ending in “_X” denote undetermined taxonomic level and were placed in the “-Unannotated” category for this study. All statistical analyses and visualizations were created in R using tidyverse, compositions, and vegan packages (Oksanen et al. 2007; Wickham 2017).

To assess ASV richness spatially, values were estimated between existing totals for ASVs detected at each station and depth assuming a consistent trend. Values were determined using a multilevel B-spline (MBA) approach for bivariate scattered data (‘mba.surf’) (Lee et al. 1997). Alpha diversity (Shannon) was determined from each sample to demonstrate relative species richness. Principal component analysis (PCA) was performed to visually represent how samples grouped (or clustered) to one another. Ahead of PCA, sequence counts per ASV were normalized across individual samples using a center log-ratio transformation. The calculated variance from the first two principal components was determined to be suitable for representation in two dimensions. K-means clustering across samples was computed using the center log-ratio transformed dataset. Nine clusters were determined to be sufficient to show the majority of sample-to-sample variability (Hartigan and Wong 1979).

Redundancy Analysis (RDA; vegan R package) was carried out to determine which environmental parameters contributed to protistan ASV composition and relative abundance. Ahead of the RDA, environmental parameters were standardized by scaling values to a zero mean (e.g., depth, temperature, salinity, etc.; a complete list of metadata are listed in Tables 1, S1) and center-log ratio normalized sequence counts were used as input. For RDA analyses at the individual taxon level, filtering by sequences per ASVs was performed on a case-by-case basis and is reported along with results. Results from the RDA were further evaluated using an ANOVA to determine the significance of each constraint. Dinoflagellate and ciliate trophic strategies were assigned using a trophic mode database from (Jones et al. 2025). To ensure accuracy, assignment between the trophic mode database and PR2 taxonomic identity results, trophic modes were assigned and curated manually.

### E. Data availability

Amplicon sequence are available at the Sequence Read Archive, under Bioproject PRJNA1495251. To reproduce data analysis, statistics, and visualizations, see code https://shu251.github.io/Gulf-2023-amplicons/.

## III. Results

### A. Northern Gulf region

Across the three transects sampled, sea surface temperatures reached a maximum of 31°C and stabilized to ≤ 5°C below 1000 m (Table 1; Figures S2-S4). Peak mixing at the MS River inflow with Northern Gulf seawater was observed at Station 1, with elevated oxygen concentrations and low salinity (∼27 ppt; Table S1; Figure S2). Transects 2 and 3 illustrated typical stratified Gulf conditions, where macronutrients nitrate, phosphate, and silicate increased with depth and nitrate decreased at depth (Figures S5-S7). Stations along Transect 1 demonstrated a similar coastal to offshore trend, but were varied at and around Station 3. Dissolved oxygen at Stations 3-4 had a subsurface increase (up to 4.5 ml L^-1^ at ∼60-100 m), while nitrite and ammonium showed subsurface drawdowns at the same depths at Station 3 (Figures S2, S5).

During the sampling period (late July-early August 2023), a Loop Current eddy named Atwater (https://www.horizonmarine.com/loop-current-eddies; separated from the Loop Current on May 18th and dissipated in October) persisted to the Southwest of the cruise transect. Stations sampled appeared to be between Atwater and a smaller warm core eddy to the Northeast, likely creating cold core eddy conditions (Figures 1B, S1). This is evidenced by the increased sea surface height (SSH) anomaly found just north of the main loop current in Figure S1B. Station 5 is the only station falling within Atwater, with all others being north of the eddy. Further, there was a localized temperature low at the surface of Station 3 (Figure S1; Supplemental Material).

The deep chlorophyll maximum (DCM; determined by a subsurface peak in fluorescence from the CTD downcast), ranged between 2-120 m, the deepest DCM was observed at stations in transect 3 (Table 1; Figures S2-S4, S8). From coastal to offshore, the salinity maxima decreased, where the lowest salinity values were observed at station 1 where the greatest influence of the Mississippi River discharge was assumed (Table 1; Figure S8). The mixed layer depth ranged between 8-30 m (Table 1; Figure S8). The oxygen minimum depth (lowest measured oxygen value during CTD cast) ranged between 46-500 m and consistently decreased from coastal to offshore stations (Table 1; Figure S8).

During the sampling period, the Mississippi River discharge averaged 263,833 cubic feet per second (cfs), reaching a peak at 290,000 cfs. Prior to this study (within 10 days of cruise start), river outflow ranged between 260,000-291,000 cfs (Figure S9). Sampling stations had between 0 to 9 oil-related structures within the 9 km buffer region (Table 1; Figure S10A). Within one week of sample collection, an active oil spill occurred within the proximity of Station 9 (Figure S10B). This study was conducted during an El Niño year (2023), which subsided into La Nina in 2024. Sea surface temperatures (SST) in July-August 2023 ranged between 28.1 and 33.1°C. During the same period in prior years (2021-2022), SST averaged 29.1 and 30.7°C. For this study, the Marine Heat Wave (MHW) Index was deemed “weak” to “borderline”. While MS River discharge, concentration of oil industry infrastructure, potential El Niño conditions, and a weak MHW index were considered in this study, they were considered outside the scope of this work due to the lack of context from seasonal or annual surveys.

### B. Summary of sequence survey

Across 12 stations, a total of 90 unique samples were recovered from depths ranging between 1.7 m to 2,827 m (Table S1). Samples were sequenced in replicate, yielding 48 million sequences.

Four samples with fewer than 40,000 sequenced reads (too few) were removed during initial sample quality control. Control samples from shipboard sampling, lab-based extraction, and PCR-based blanks were also sequenced in order to identify and remove 766 ASVs as potential contaminants. To further quality control data for this study, ASVs assigned to “Unassigned”, “Bacteria”, or “Eukaryota:nucl” were removed, yielding 38.4 million sequences and 38,262 ASVs. ASVs from duplicate samples, when possible, were averaged by taking the mean sequence count and all ASVs were required to have ≥ 100 sequences across the whole data set. Following these quality control filtering steps, the dataset consisted of 10,563 ASVs and 21 million sequences.

### C. Microbial biodiversity across environmental gradients in the Northern Gulf

The total number of ASVs ranged between 400 to 2000 for samples at the surface and throughout the water column (Figure 2, Table S1). There were two surface ASV maxima observed at Stations 7 and 9 (up to 2000 ASVs each; Figure 2A), which aligned with elevated species richness primarily within the upper 200 m (Figures 2 and 3). Alpha diversity at the surface and upper water column showed the greatest amount of variability (Shannon diversity values of 2-6) and included the lowest alpha diversity metrics (Figure 3A). Ordination analysis revealed samples to group largely by depth in the water column (Figure 3B). Samples at the surface and subsurface up to ∼270 m varied in ordination space, with two main groups composed of mostly subsurface and then mostly surface samples (Figure 3B). Deeper, offshore samples (> 500 m) formed a tighter group relative to samples from all other depths (Figure 3B).

**Figure 2.**
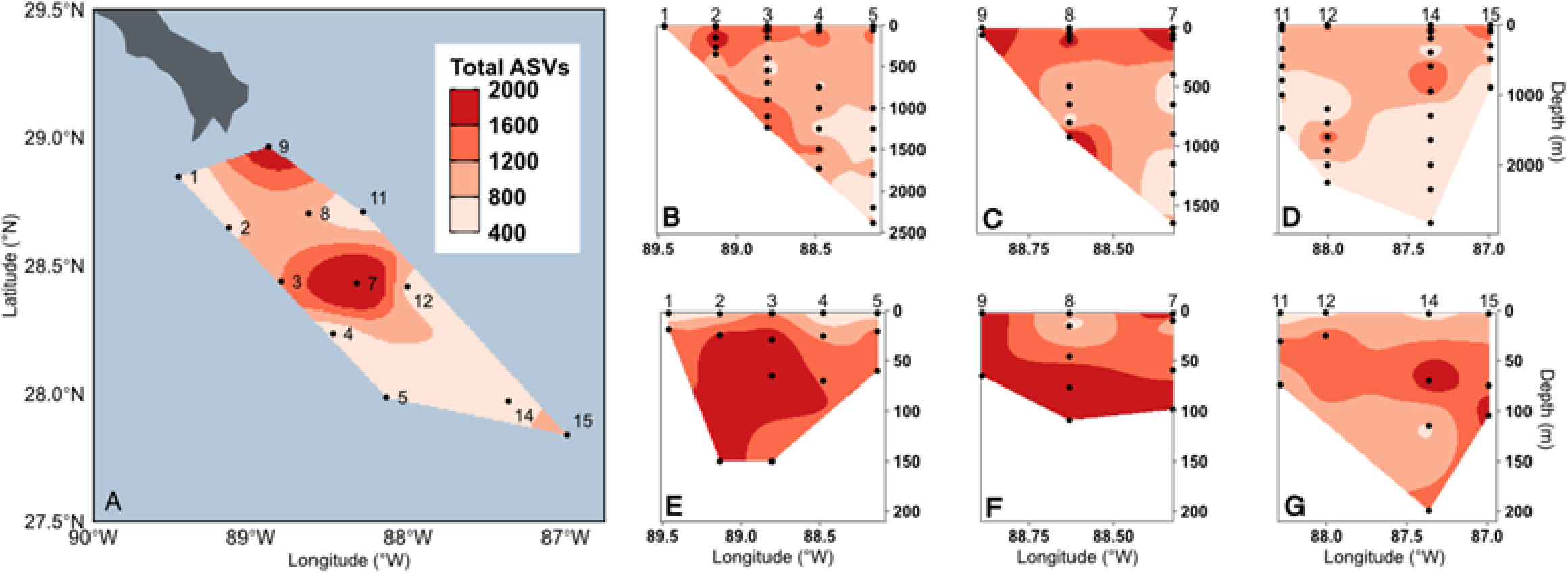
Total number of Amplicon Sequence Variants (ASVs; heat map) recovered from each surface station, the complete vertical profile from transects 1-3 (B-D), and the upper 200 m of the water column at transects 1-3 (E-G). Mapping of ASVs spatially was interpolated by determining using a multilevel B-spline between points.

**Figure 3.**
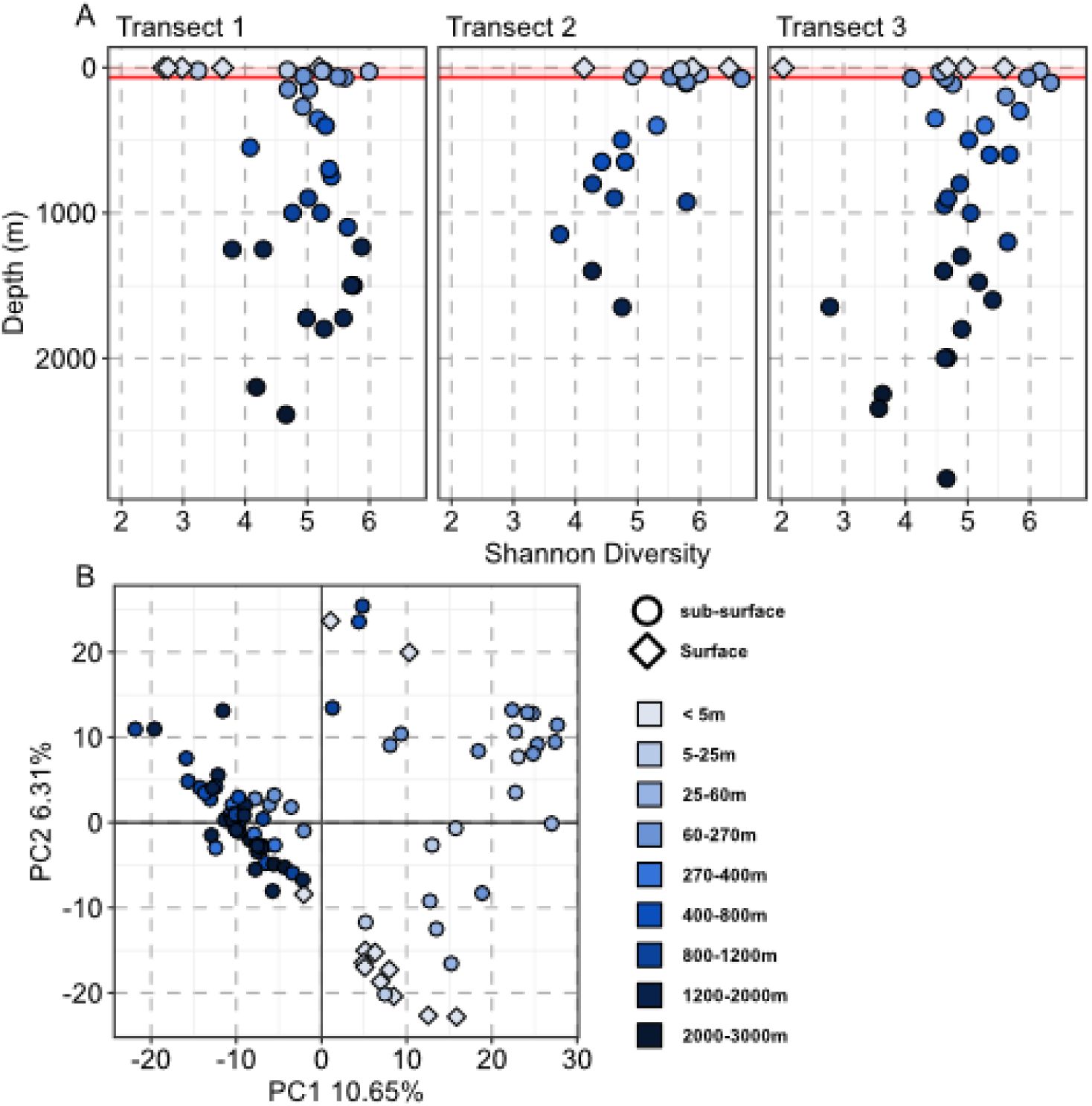
(A) Shannon diversity (y-axis) for each sample by station and transect (x-axis). Similar to (B), colors denote depth bin (m) and symbol indicates if the sample is from the surface (<5m) or below the surface.(B) Principal component analysis (PCA) for all samples based on approximate depth (meters; color of symbols). Symbol indicates if the sample is from the surface (<5m) or below the surface. Data point colors denote depth bin (m) and symbol indicates if sample is from the surface (<5m) or below the surface.

K-means clustering resulted in 9 sample groupings that were structured by depth, water masses, proximity to coastline, nutrient profiles, and protistan taxonomic composition (Figure 4). Clusters 1, 4, 7, and 8 primarily included the upper water column samples and were attributed to Caribbean Surface Water and Gulf Common Water. The majority of samples within Clusters 1, 4, 7, and 8 originated from above the mixed layer depth (MLD), DCM, oxygen minimum, and the salinity maximum features and included higher relative abundances of haptista, archaeplastida, and ciliates relative to other taxa (Figure 4). Clusters 2, 3, 6, and 9 included the majority of deep seawater samples, and included a mixture of Tropical Atlantic Central Water (Figure 4). Samples from below the MLD, DCM, oxygen minimum, and salinity maximum were primarily assigned to Clusters 2, 3, 6, and 9. Broadly, nutrient profiles, especially for oxygen, nitrite, nitrate, ammonium, and phosphate were elevated within Clusters 2, 3, 6, 9 (Figure 4). Coastal surface samples were made up of Clusters 2 and 5, with Cluster 5 encompassing the highest total number of surface samples (< 5 m).

**Figure 4.**
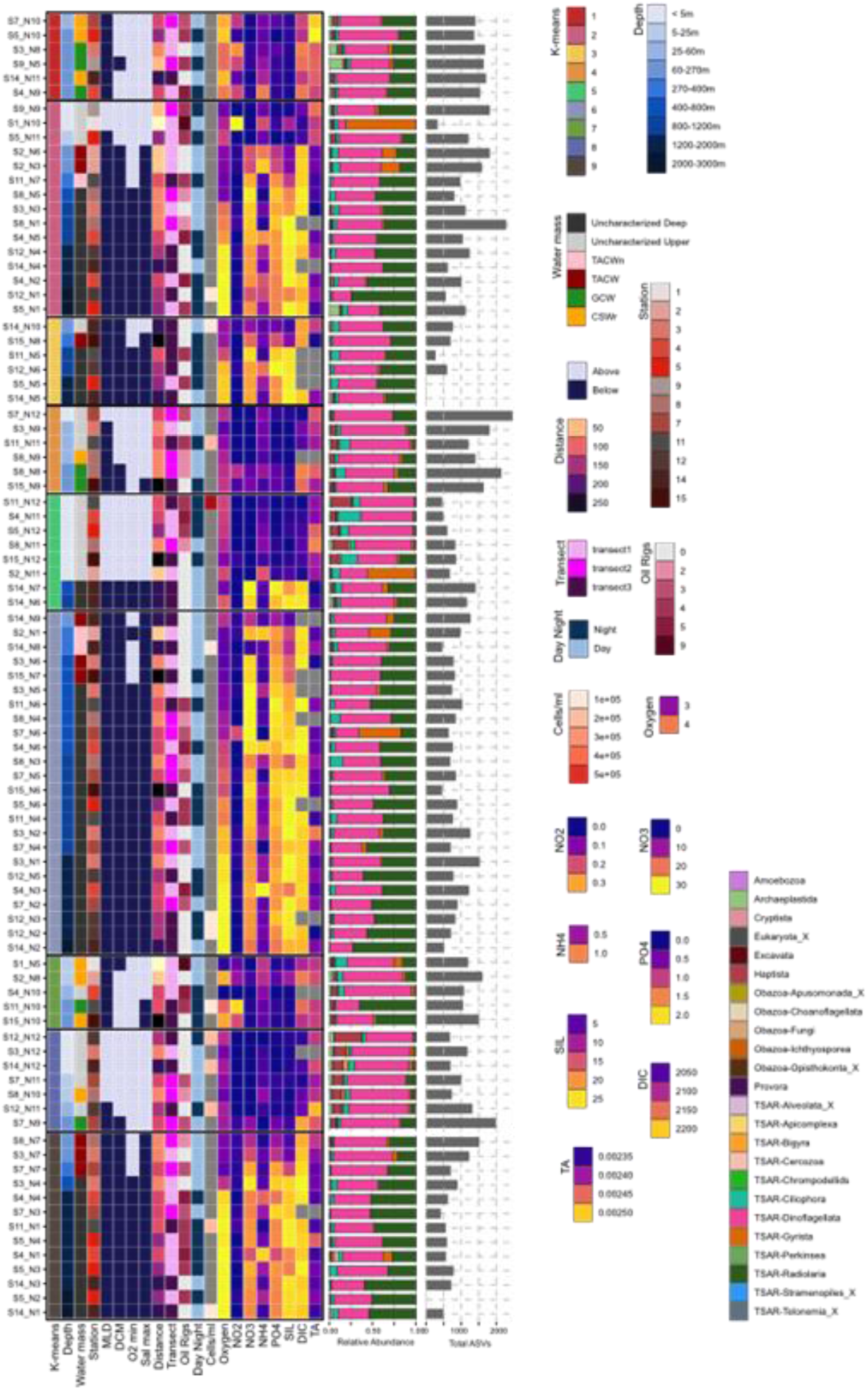
Compilation dendrogram for all samples in this study. Samples (Station and Niskin ID labeled) are listed based on the k-means cluster membership (n=9; furthest left). From left to right, heat map colors indicate depth range in the water column (m), closest associated water mass, station number, if the sample is situated above or below the mixed layer depth (MLD), deep chlorophyll maximum (DCM), oxygen minimum layer (O2 min), or the salinity maximum (Sal max), the approximate distance to the coast (km), transect (1, 2, or 3), total number of oil rigs within a 10 km radius, if the sample was collected during the day or night, microbial cell concentration, oxygen, nitrite, nitrate, ammonium, phosphate, silicate, dissolved inorganic carbon (DIC), and total alkalinity (TA). On the right, the relative sequence abundance is shown as a bar plot, where colors denote major protistan taxonomic groups and the furthest right bar plot reports the total number of recovered ASVs from each sample.

RDA results demonstrated the relative importance of how specific environmental conditions and sample types shape protistan biodiversity. Broadly, environmental parameters constrained a significant portion of protistan community composition, although the identity of the significant drivers varied with depth (Table 2). Altogether, 44% of the microeukaryotic community composition was explained by the available metadata: (in order of significance) water mass, depth of the DCM, sampling depth, and distance to the coast (Table 2). Depending on the taxonomic group, the depth of the DCM or MLD, and concentrations of nitrite, nitrate, phosphate, and the ratio of silicate:nitrate were also influential (Table 2). When samples were separated by into layers above and below the DCM, the raw constrained fraction was higher above (84%) than below (37%; this different largely reflects the smaller number of samples retained above the DCM, and after adjusting for sampling effort the two layers explained comparable variation (adjusted R^2^ = 0.09 and 0.12, respectively). Significant drivers differed between the layers; below the DCM, sampling depth, distance to the coast, nitrate, and phosphate each structured community compositions, whereas above the DCM none of these parameters were significant (Table 2).

**Table 2.**
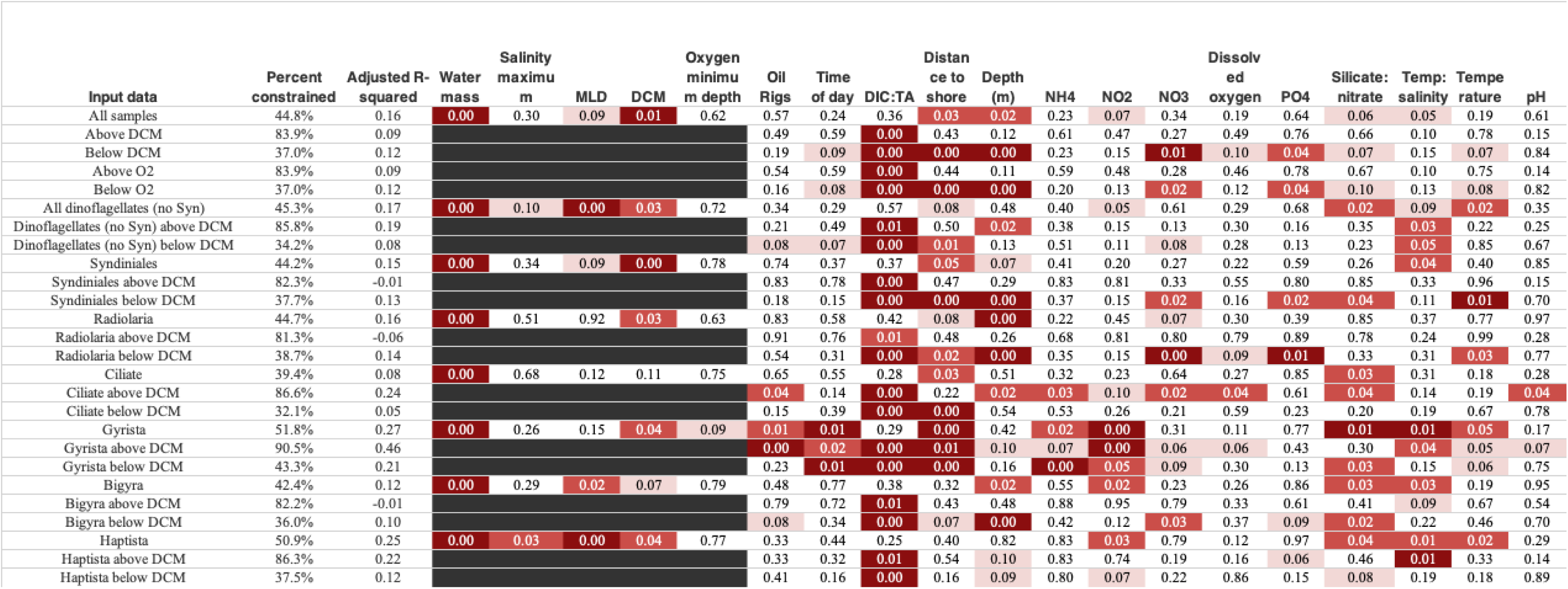
RDA analysis results demonstrate how varying environmental parameters explain protistan biodiversity across stations, depth, and metadata. Input data on the left-most column highlights the subset of the microbial community examined for each test (by row). Percent constrained represents the portion of the microeukaryotic community that can be explained by the input variables. Values below metadata report significance for the input data and are color-coded for significance level.

### D. Taxon-specific patterns

Surface and depth-integrated ASV richness by taxa (Figures S12-S15) somewhat mirrored the total community (Figure 4). Broadly, dinoflagellate sequence relative abundance often exceeded 50% of the recovered sequences across each sample; exceptions to this included samples from deep depths, where there was a higher relative abundance of radiolaria sequences observed.

Several samples also had higher relative abundances of gyrista at depth (Figure 4). Within dinoflagellates, the majority were found to belong to the primarily parasitic class Syndiniales, which was dominated by Dino-group-I or Dino-group-II across all depths (Figures 5A and S12B). Syndiniales ASV richness was elevated at several surface stations, most notably station 7 (Figure S12B). Syndiniales ASV distribution below the DCM was found to be constrained by nitrate, oxygen, and distance from the coastline (Table 2). Following Syndiniales, the

**Figure 5.**
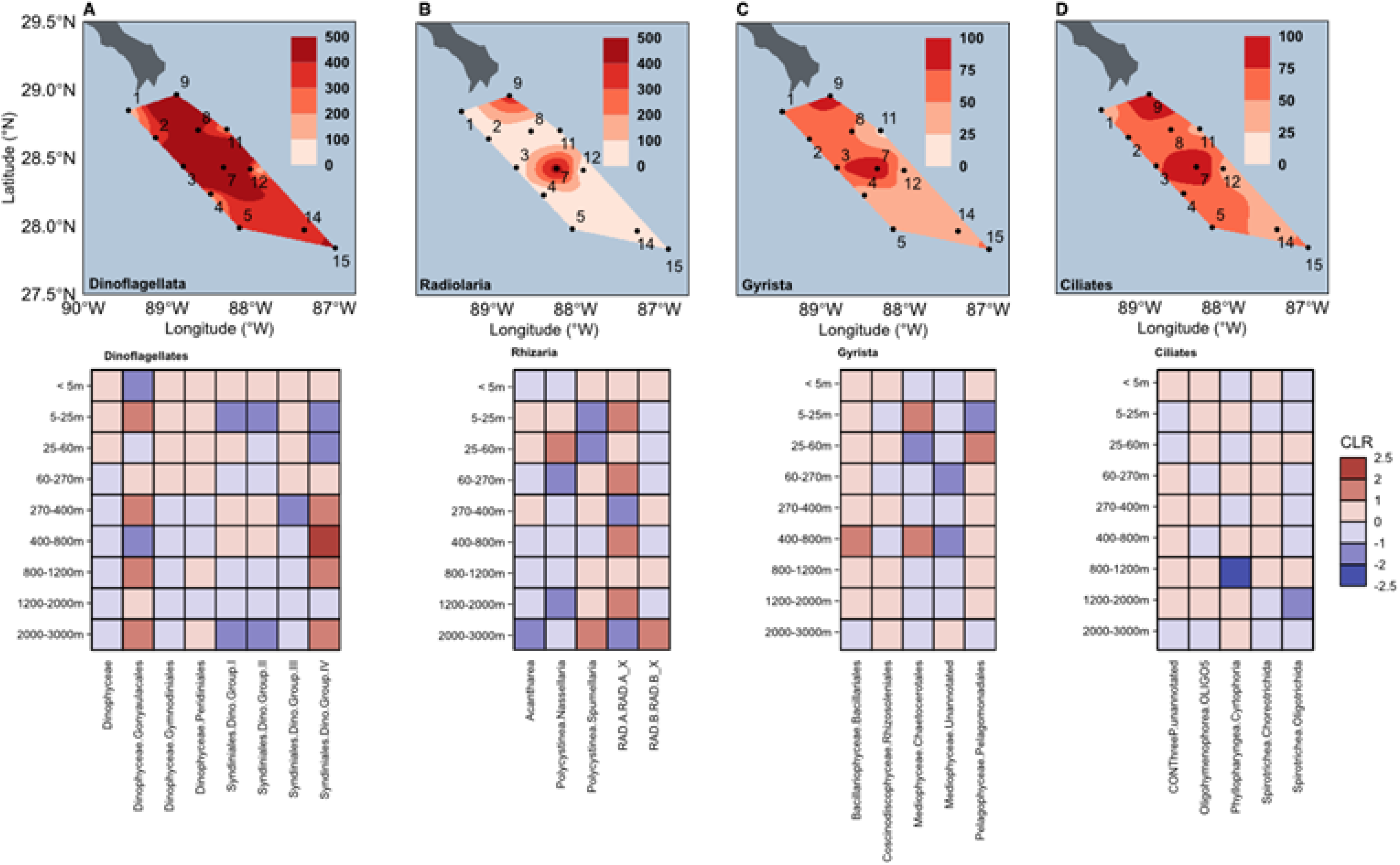
Relative sequence abundance for (A) dinoflagellates, (B) radiolaria, (C) gyrista, and (D) ciliates. For each panel, samples were combined based on membership to coastal (top left; stations 1, 2, 3, 8, and 9) versus offshore stations (bottom left) or above 200 m (shallow; top right) versus at and below 200 m (deep; bottom right). Each panel represents the composition of total ASVs belonging to phyla or classes within each major taxonomic group.

Dinophyceae class was the primary dinoflagellate recovered. While relative sequence abundances varied with respect to proximity to shore at the surface, *Gymnodiniales* was detected at all depths and stations, while the *Gonyaulacales* was not found at depth (Figure 5A).

Depending on location and depth, ASVs identified as Radiolaria or gyrista (stramenopiles) were the next most numerous taxonomic group. Radiolaria were primarily made up of *Spumellaria* (*Polycystinea*), Acantharea, and the radiolarian environmental clades RAD-A, −B, and -C across all samples (Figure 5B). Coastal and upper water column samples were characterized with higher relative abundances of the *Polycystinea* group *Nassellaria* (Figure 5B). Environmental clades RAD-A, -B, and -C had distinct distributions with respect to depth: RAD-A was found throughout the coastal and upper water column samples (<200 m), with RAD-B primarily restricted to deeper depths (Figures 5B and S13). The maximum depth of the station sampled, nutrient profiles, and oxygen were all found to significantly contribute to radiolaria ASV distribution (Table 2).

ASVs belonging to Gyrista included members of the *Bacillariophyceae, Chrysophyceae, Coscinodiscophyceae, Dictyochophyceae, Mediophyceae*, and *Pelagophyceae* (Figure 5C). The three primary Gyrista classes found were Mediophyceae, Coscinodiscophyceae, and Bacillariophyceae. Higher sequence abundances for Gyrista were generally found at coastal, shallow sites and were accompanied by increases in total Gyrista ASVs (Figure S14). Surface stations situated closest to the coast, specifically Stations 1 and 2, were primarily comprised of ASVs identified as unannotated or *Hemiaulales* (*Mediophyceae*) and *Rhizosolenia* (*Coscinodiscophyceae*). The composition of Gyrista shifted immediately offshore, where there was a relative increase in Chrysophyceae (Stations 3 and 8), specifically Ochromonadales and Paraphysomonas (Figure 5C). All macronutrients played a role in shaping the Gyrista community composition, except for phosphate (Table 2). At subsurface depths, both relative sequence abundances and total gyrista ASVs were lower (Figure 4).

While ciliates were not numerically dominant throughout the sampling sites, they were present in all samples and displayed depth-specific patterns. The ciliate community was primarily made up of Spirotrichea groups Choreotrichia and Oligotrichia (Figure 5D). Relative sequence abundances for Choreotrichia were higher in deeper, offshore samples (Figure 5D). The relative abundance of Phyllopharyngea was higher across all upper water column samples, while at depth, the relative abundance of scuticociliates, apostomatia, and suctoria increased (Figure 5D). Ciliates found above vs. below the DCM were subject to different parameters that shaped biological diversity (Table 2; RDA).

### E. Functional trait assignment for dinoflagellates and ciliates

Over half of the dinoflagellate ASVs (67%) and roughly a third of the ciliate ASVs (29%) were assigned trophic modes using the Jones et al. 2025 database. Among dinoflagellates, there were autotrophic members of the *Gonyaulacales*, strict heterotrophic *Gymnodiniales*, and almost 100 constitutive mixotrophs from the *Gymnodiniales*, *Gonyaulacales*, and *Prorocentrales* (Figure 6). Out of approximately 5500 dinoflagellate ASVs, over 3600 belonged to Syndiniales, which were assigned as parasitic and found at all depths and samples in this study (Figures 6, S16).

**Figure 6.**
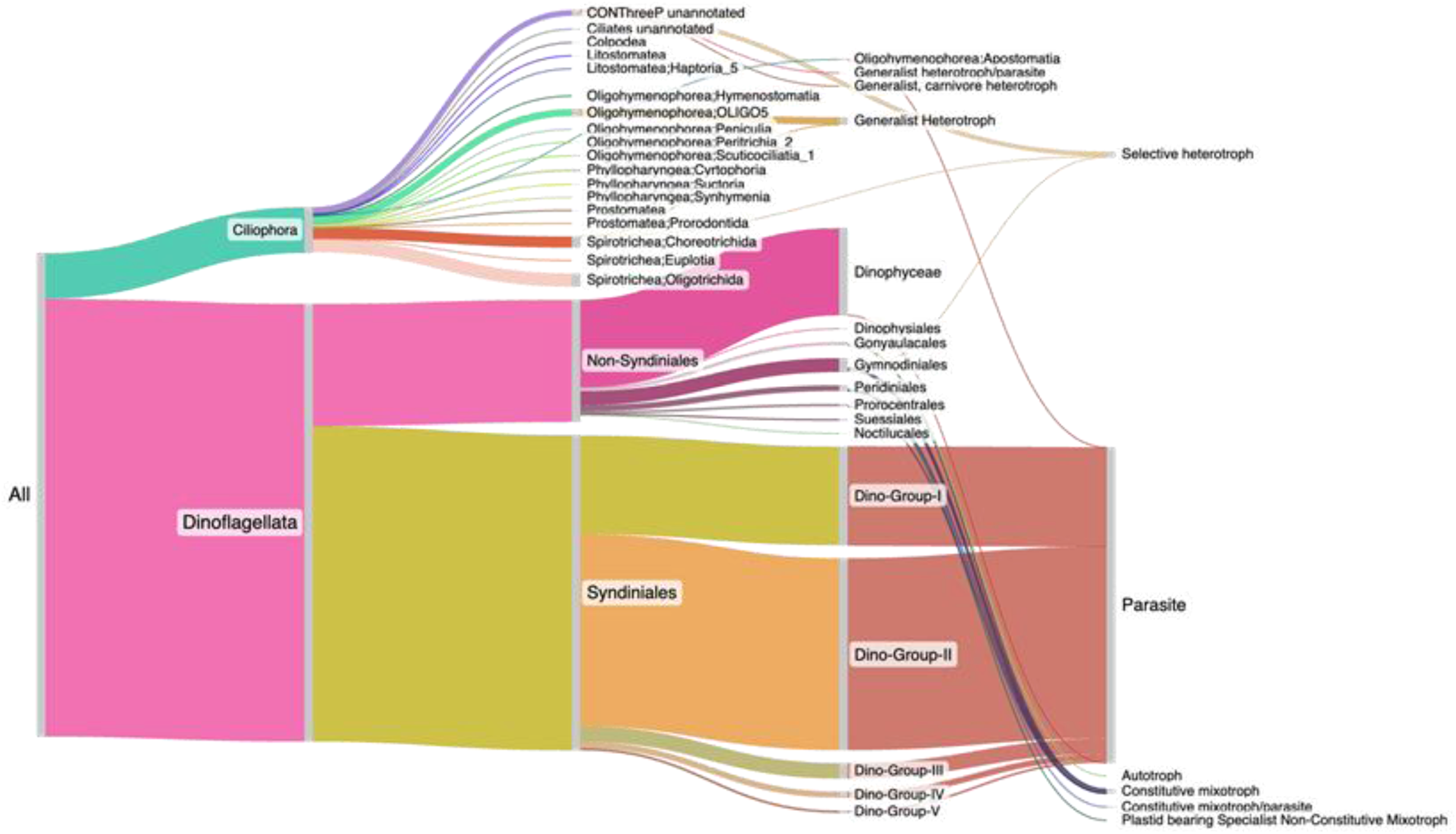
Sankey diagram illustrating the distribution of ciliate and dinoflagellate ASVs identified by trophic mode category.

Following Syndiniales parasites, constitutive mixotrophs were the most abundant trophic strategy represented by dinoflagellates. Dinoflagellate mixotrophs and heterotrophs were found across all depths, with a general increase in ASVs identified as putative constitutive mixotrophs (Figure S16, Table S2). While fewer ciliate ASVs were assigned trophic strategies compared to dinoflagellates, members of the CONthreeP group and Choreotrichs were labeled as selective heterotrophs, while other Spirotrichea and Oligohymenophorea were identified as generalist heterotrophs (Figures 6, S16, Table S2). The highest number of both selective and generalist heterotrophic ciliates was found within the 60-270 m range (Figure S16).

## IV. Discussion

The Northern Gulf represents a natural incubator to assess how a range of environmental forcing factors may select for marine microbial species. Details of phytoplankton community dynamics (species composition and structure) in the Gulf rely largely on satellite-derived observations, leaving observations of subsurface microbial communities unconstrained (E Selph et al. 2022). Depth-related shifts in the microeukaryotic community have been documented (Landry et al. 2022; Linacre et al. 2025), but linking them to large-scale environmental gradients remains under-studied. In the Northern Gulf, we found microeukaryotic biodiversity to be dictated by the interplay of the Mississippi riverine input, Loop Current eddy shedding, and resulting mesoscale features. Our findings highlight how depth, distance to coastline, nutrient profiles, and submesoscale features drive microeukaryotic biodiversity, reinforcing protists as ideal ecosystem indicators.

### A. Protistan biodiversity shaped by depth & proximity to coastline

The significance of water mass, DIC:TA, distance to the coast, and depth in the water column played the largest role in structuring protistan biodiversity; this suggests several co-factors contribute to overall community structure. Our survey revealed that protistan communities below the DCM were more likely to be constrained by macronutrient concentration (nitrate and phosphate) and distance from the coastline, whereas above the DCM, these gradients were not significant (Table 2). Taken together, the DIC:TA ratio was significant within each depth layer, while water mass and depth were significant when considering the whole dataset; this demonstrates that water mass and DIC are likely covariants. DIC will be drawn down in the upper water column due to carbon fixation processes by the resident microbial community (Table 2), which are highly dependent on the influence of water mass mixing and coastal runoff (Cai et al. 2011; Wang et al. 2013; Hu et al. 2018). The ratio of silicate to nitrate played a significant role in shaping the phytoplankton communities, specifically dinoflagellates, gyrista, bigyra, and haptista (Table 2). The ratio of silicate to nitrate is recognized as a core variable structuring coastal phytoplankton communities: diatoms become silica limited as the ratio approaches 1:1 and the community typically shifts towards a higher abundance of non-siliceous flagellates (Officer and Ryther 1980; Turner et al. 1998) and has been documented within the MS River plume (Krause et al. 2023). Since silica and nitrate have distinct biological sinks and depth profiles in the water column, the ratio of Si:N may serve as a sensitive predictor of phytoplankton composition. This biological coupling has significant implications for regional biogeochemistry: the protistan assemblage favored by a particular Si:N ratio determines the ultimate fate of sequestered carbon. A community dominated by silica-ballasted diatoms will have a high carbon export efficiency, due to rapid sinking. Alternatively, a non-siliceous flagellate and/or dinoflagellate community is more likely to increase remineralization processes in the upper water column (Tréguer et al. 2018). The simultaneous tracking of silicate and nitrate connects microeukaryotic biodiversity to Gulf carbon cycling, suggesting that the Si:N ratio may indicate how future variations in Mississippi River nutrient loading could restructure Northern Gulf food webs.

The observed shifts from diatoms within nearshore stations to dinoflagellates at surface offshore stations and rhizaria at depth in the Northern Gulf may have implications for the coupling of carbon export between the upper water column and deep sea, although carbon export was not directly measured in this study. Both diatoms and rhizaria are known to make up a substantial fraction of sinking particle flux (Boeuf et al. 2019; Beatty et al. 2025). In a more diatom dominated community, rapidly sinking diatoms are known to contribute to increased carbon export efficiency (Tréguer et al. 2018). Classes within the dinoflagellates, rhizaria, and ciliates also demonstrated depth partitioning, which can be attributed to presumed trophic mode or symbiotic partnerships with other microbes. We examine these groups, dinoflagellates, diatoms, rhizaria, and ciliates, below as ecologically important microorganisms at the base of the Gulf-based food web.

Rhizaria displayed the most pronounced depth partition across the three groups: within the Rhizaria sequences recovered, both acantharia and spumellaria were the most numerous. The morphologically-elusive environmental clades RAD-A, -B, and -C are widely distributed across the world’s ocean (Mars Brisbin et al. 2020; Biard 2022; Sandin et al. 2025) and displayed depth-specific niche partitioning in this study. Prior work in the Gulf has noted RAD-A, -B, and Spumellaria to be negatively correlated with temperature and positively correlated to nitrate (Sidón-Ceseña et al. 2025) and RAD-A has previously been linked to anticyclonic eddy activity. We found both RAD-A and -B to be enriched at eddy edge stations (e.g., 7 and 8; Figure S13) and ASVs identified as RAD-B to consistently increase at greater depths; the latter of which has been observed in prior sequence-based surveys (Giner et al. 2019).

Similar to prior work on protistan biodiversity in the Gulf, dinoflagellates were the most abundant 18S rRNA gene signature recovered (Figure 4). While recovery of dinoflagellate ASVs can be misleading due to the wide range in gene copy number (Liu et al. 2021), we used both ASV presence/absence and center normalized methods to evaluate dinoflagellate distribution.

Further, prior work using quantitative cell counts or imaging (e.g., IFCB), and HPLC pigment analysis has found dinoflagellates to make up a substantial amount of the microbial carbon biomass in the Gulf (Linacre et al. 2025; Sidón-Ceseña et al. 2025). Investigating the relative dominance of specific dinoflagellate species is necessary, as they encompass a dynamic range of nutritional strategies (feed modes) that have varying impact on primary productivity, food web efficiency, and carbon export (Bi et al. 2021).

The exclusively parasitic Syndiniales clades made up a substantial proportion of the recovered dinoflagellate sequences and displayed depth-specific patterns across the Northern Gulf (Figure 5A, S12). At the Bermuda Atlantic Time-series (BATs), depth-specific trends in Syndiniales clades were found to correspond to light and oxygen availability (Anderson et al., 2024). These patterns and overall Syndiniales biogeography in the ocean is thought to be driven by host composition, which include metazoa (e.g., copepods, larvae), dinoflagellates, haptophytes, or ciliates (Chambouvet et al. 2008; Lima-Mendez et al. 2015). Dino-Group I was previously shown to have a higher relative sequence abundance within dark, oxygen-depleted conditions, while Dino-Group II was found throughout the water column. Dino-Group I is also known to be associated with rhizarian hosts, specifically Polycystinea, acantharea, and RAD-B, which aligns with our observations of increased rhizarian genetic signatures at depth (Figures 4, 5). However, the confirmation of specific host-parasite relationships and parasitic infection cycle *in situ* remains a substantial challenge in marine microbial ecology. Regardless, environmental surveys that constrain the distribution and extent of Syndiniales groups bring us closer to incorporating parasitic dynamics into marine food web models (Jephcott et al. 2016; Ollison et al. 2021; Anderson et al. 2024).

Non-Syndiniales dinoflagellate sequences included members of the Dinophyceae, such as Gonyaulacales, Gymnodiniales, and Peridiniales (Figure 5A). With half of these assigned confirmed trophic strategies using the Jones et al. (2025) database, we can begin to see how specific traits may be selected for *in situ*. Overall, non-constitutive mixotrophic dinoflagellate taxa had more restricted spatial distributions. At depth, ASVs identified as Peridiniales and Noctilucales, which are symbiotic non-constitutive mixotrophs, are likely contributing to the microbial food web as heterotrophs. Gymnodiniales’ success in the Gulf as mixotrophs, heterotrophs, and parasites is supported by these findings and similar to other work in the Gulf (Sidón-Ceseña et al. 2025). Our findings suggest that mixotrophy is likely the “norm” for Gulf-based dinoflagellates and contributes to their ecological success in this dynamic region.

Ciliates showed a comparable depth structure among the heterotrophic taxa recovered. Ciliates were present across all depths and stations sampled and, while not numerically dominant (with respect to relative sequence abundance), were observed to exhibit depth-specific distributional patterns that are consistent with their presumed trophic modes (Mitra et al. 2023; Jones et al. 2025). Ciliates in the Northern Gulf were primarily made up of Spirotrichea, specifically Choreotrichia and Oligotrichia (Figure 5D), which is similar to prior descriptions of ciliates in the Gulf (Snyder et al. 2021) and other highly stratified regions (Johansson 2004). Choreotrichs and oligotrich ciliates are widely recognized as the primary ciliate consumers of nanoplankton and small phytoplankton, and thus represent a critical trophic link between primary producers and higher trophic levels (Santoferrara et al. 2023). Below the DCM and oxycline, relative abundances of scuticociliates, apostomatia, suctoria increased relative to the upper water column depths (Figure 5D). This trend is similar to the previously observed euphotic to subeuphotic transition in mixotrophic, algivorous ciliates to primarily bacterivorous and parasitic ciliate taxa below the reach of sunlight (Santoferrara et al. 2023).

Together, assignment of putative feeding modes for both dinoflagellates and ciliates demonstrated the impact of the presence of these ecologically important species relies on how they may be interacting with their environment and other microorganisms. The impact that mixotrophy has on food webs has been documented in global biogeochemical models and laboratory-based experiments (Ward and Follows 2016; Millette et al. 2023). Ciliates assigned as either selective or generalist heterotrophs will have implications for how carbon may be assimilated and transferred from primary producers to higher trophic levels. Linking documented prey preferences or individual species with specialist versus generalist grazing strategies (Mitra et al. 2023; Jones et al. 2025) has broader implications for food web dynamics. More specifically, selective heterotrophic ciliates (e.g., Choreotrichia) and generalist ciliate species (e.g, oligohymenophora) will have distinct efficiencies of carbon transfer to higher trophic levels and relative contributions to DOM (Maiti et al. 2016). Altogether, this work highlights the need to capture *in situ* contributions to the food web among mixotrophs and heterotrophs, as, together, cell physiology and environmental conditions will dictate what trophic strategy they ultimately exhibit.

### B. Symbiotic partnerships at the river-seawater interface & eddy edge

Diatoms are known to outcompete other phytoplankton in turbid, high nutrient conditions (Fawcett and Ward 2011; Edwards et al. 2015) and have previously been the dominant member of the phytoplankton community where the MS River input mixes with the Gulf (Chakraborty and Lohrenz 2015; Chakraborty et al. 2017; Anglès et al. 2019; Krause et al. 2023). Estuarine and shelf regions in the Northern Gulf have specifically been dominated by *Asterionellopsis*, *Chaetoceros*, *Skeletonema*, *Pseudo-nitzschia*, and Thalassiosirales similar to (Anglès et al., 2019). In this study, stations most heavily impacted by the MS River captured a surface layer made up of over 50% diatoms associated with the classes *Mediophyceae* and *Coscinodiscophyceae (Figure 5C*; Table 2). Stations 1 and 2 were specifically dominated by *Hemiaulus (Mediophyceae)* and *Rhizosoleniales* (*Coscinodiscophyceae*; Figure S14). *Hemiaulus and Rhizosoleniales are* diatom-diazotrophs that harbor nitrogen-fixing cyanobacteria (Karl et al. 2012; Kemp and Villareal 2018; Pyle et al. 2020; Flores et al. 2022). We hypothesize that as Mississippi River discharge is lower and the springtime influx of nutrients has been drawn down in late summer (July to August; period of our study), diatoms associated with nitrogen-fixing cyanobacteria (*Richelia intracellularis*) increase in abundance. Because our sampling captured a single week, we present this as a hypothesis rather than observed succession: seasonal shifts in diatom species within the Northern Gulf may have varying contributions to carbon and nitrogen fixation, particularly in providing biologically available nitrogen to the region.

Further, Linacre et al. (2025) found evidence of the haptophyte prymnesiophyte species that host the diazotrophic cyanobacterium UCYN-A. This observation was further supported from a paired study that detected an elevated contribution to new nitrogen within the Loop Current Eddy, relative to outside the eddy (Herzka et al. 2024; Linacre et al. 2025). Similar to our findings, the putative host relationship with N_2_-fixing symbionts is inferred for prymnesiophytes, where there was an increase in ASV richness for prymnesiophytes at Stations 3, 8, and 12 (Figure S12A). These taxon-specific routes of potential N_2_ fixation throughout the Northern Gulf warrant additional study, especially paired with targeted rate measurements, confirmation of host-symbiont identity, and temporal resolution to address potential seasonal trends.

### C. Linking shifts in biodiversity to mesoscale features

Offshore, an eddy shed from the Loop Current (named Atwater) remained to the south of our study site and a second began to form directly to the east; together, these features created a cold core eddy-like condition situated around Station 3 (Figures 1, S1, Supplemental Material). We detected increased protistan species richness at the edge of this eddy, localized to the upper water column at Station 7 (Figures 2A and 3A). This has been similarly observed within the North Pacific Subtropical Gyre (NPSG), where increased phytoplankton species richness was found within eddies and along the eddy edge where nutrients are redistributed (Jones-Kellett et al. 2026) and in the Southern Indian Ocean and South China Sea, where mesoscale features were linked to an increase in microeukaryotic biodiversity (Wu et al. 2015; Sturm et al. 2023). The biological impact of these physical features can also be seen within prymnesiophyceae, dinoflagellates, rhizaria, gyrista, and ciliates (Figures S12-S15) and reinforced by elevated subsurface dissolved oxygen (Figure S2D) and nitrate (Figure S5) concentrations. Mesoscale circulation features are known to deliver subsurface nutrients to the upper water column and subsequently cause increased phytoplankton primary productivity, especially at an eddy edge (Biggs 1992). Edges of eddies can entrain either coastal MS River plume water and/or deep, nutrient-rich seawater and form a frontal boundary enriched in microbial biomass and activity (Damien et al. 2021; Linacre et al. 2025). Differences in eddy polarity have also exhibited disproportionate contributions to sinking particle flux by microeukaryotes; in Beatty et al. (2025), while the protistan community composition across cyclonic vs. anticyclonic eddies was not significant, the relative contribution of siliceous organisms within captured flux material was linked to the cyclonic eddy polarity. Due to our observed shifts in microeukaryotic biodiversity across mesoscale features, we suspect that the contribution of microeukaryotes to particle flux is dependent on physical forcing factors in the Northern Gulf.

Mesoscale features can restructure protistan trophic modes, independent of taxonomic identity. In the oligotrophic North Pacific Subtropical Gyre (NPSG), cyclonic eddies subject to upwelling selected for protists known to have heterotrophic traits, while anticyclonic eddies favored transcript signatures more closely associated with photoautotrophic modes of nutrition (Harke et al. 2021; Gleich et al. 2024). Observed shifts in trophic mode persisted despite the taxonomic composition remaining similar, indicating that the mesoscale forcing factors influenced the physiological response of the resident protistan community (Harke et al. 2021). Prior work in the Northern Gulf found cyclonic and anticyclonic mesoscale features to impact plankton assemblages (<10 µm), specifically noting eddy boundaries to have distinct populations of autotrophs and heterotrophs (Williams et al. 2015). Consistent with this trend, we recovered elevated species richness and a distinct protistan community signature at both station 3 and 7 (Figures 2, 3A). These samples were characterized by higher relative abundances of ASVs identified as haptista, a group that includes many mixotrophic species, and likely drove the k-mean cluster group 2 formation (Figures 4 and S11; Table S1). Station 7 was also characterized by a relative increase in the silicoflagellate Dictyochophyceae and picoeukaryotic Pelagophyceae, which are known to be more successful in oligotrophic conditions. *Pelagomonas* was found primarily in subsurface depths, aligning with prior work showing Pelagomonas to prefer lower light intensities (Rii et al. 2016; Sidón-Ceseña et al. 2025). Overall, we found that the introduction of nutrients from eddy-driven mixing paired with a sharp light vertical gradient can create heterogeneous conditions that will select for specific protistan trophic modes, specifically for smaller mixotrophic protistan species rather than diatoms.

### D. Summary

The Gulf’s recovery capacity can be determined by evaluating if the ecosystem is equipped with enough microbial diversity for it to return to equilibrium after a hazardous event. The Northern Gulf region is changing and the roles protists play in the food web have the potential to greatly impact how carbon is shuttled and transformed in the food web. We show here how the protistan community is influenced by regional physical oceanography and environmental changes.

Molecular signatures of marine protistan biodiversity (derived from 18S rRNA gene metabarcoding) reflected regional oceanographic conditions. Our results expand upon prior work to demonstrate that within the river-influenced, productive margin system, eddy-driven nutrient dynamics interact with riverine inputs to produce a gradient of varied protistan trophic strategies. Continued, multi-depth monitoring of microbial biodiversity across the coastal to offshore gradient in the Northern Gulf represents an invaluable baseline for understanding how microeukarytic communities will respond to anomalous environmental changes (e.g., marine heat waves, oil spills, or El Niño) and for assessing the broader resilience of the Northern Gulf ecosystem.

## Acknowledgements

Authors would like to acknowledge the Captain and crew of the RV Point Sur, the science party of the GRADients 2023 research cruise, and Shari Yvon-Lewis. Student participation was made possible by the Undergraduate Research Program at Texas A&M University.

Sequence libraries were prepared and sequenced at the Georgia Genomics and Bioinformatics Core acknowledgement: GGBC, UG Athens, GA, RRID:SCR_010994.

---

**Table S1.** Complete sample information, environmental conditions, and raw sequence file information for the study. From left to right, columns report the station sampled (see map Figure 1A), niskin bottle sampled during seawater collection, depth of sample (meters), latitude and longitude, date and time of collection (UTC), if collection was done during night or day, transect (1-3), distance (m) from Mississippi River outflow, binned depth for sample (m), number of oil rigs within a 9 km radius of sampling location, depth of the mixed layer (MLD in meters), depth of the maximum salinity value (m), depth of the minimum dissolved oxygen value (m), deep chlorophyll maximum depth (m), coastal vs. offshore classification, membership to the k-means clusters, dissolved inorganic carbon (µM), NH_4_^+^ (µmol L^-1^), NO_2_ (µmol L^-1^), NO_3_^-^ (µmol L^-1^), dissolved oxygen (ml L^-1^), PO_4_^-3^ (µmol L^-1^), Si(OH)^4^ (µmol L^-1^), salinity (ppt), Total alkalinity (TA, µmol L^-1^), temperature (°C), pH, prokaryote cell counts (cells ml^-1^), names of the forward and reserve reads (fastq files), total number of recovered ASVs, and sequences .

**Table S2**. List of ASVs assigned trophic mode using Jones et al. 2025.

**Figure S1**. (A) Regional map showing the southeast United States and Mexico. The rectangle outlines the study area shown in B and C. (B) Sea surface height (SSH; m) of study area. Data points show the location of stations 1-5. (C) Sea surface temperature (°C) during the study period. Data points show the location of stations 1-5.

**Figure S2-S4.** Water column parameters for Transect 1, 2, and 3. Stations are noted at the top of each panel with a vertical line through the profile and the potential density is mapped on top of each profile. (A) Temperature (°C) from the surface to deepest depth, and (B) from the surface to 500 m. (C) Oxygen (ml/L) from the surface to deepest depth and (D) from the surface to 500 m.

(D) Salinity (PSU) for the full vertical profile and (F) up to 500 m. (G) Fluorescence (mg m^-3^) throughout the full profile and (H) up to 500 m.

**Figure S5-S7**. Nutrient profiles for Transects 1, 2, and 3. ODV interpolation derived from discrete samples (white symbols) from each station (numeric at top of each panel). All units are in µmol L^-1^ and the full profile (surface to deepest depth) is shown on the left-hand side panels and paired with the same profile up to 500 m on the right-hand side. Profiles are shown for (A-B) nitrite, NO_2_, (C-D) nitrate, NO_3_, (E-F) ammonium, NH_4_, (G-H) phosphate, PO_4_ (I-J) and silicic acid (SIL).

**Figure S8.** Depth features across each transect (panels from left to right) and station (x-axis bottom labels). Symbols and colors denote the mixed layer depth (MLD), deep chlorophyll maxima (DCM), depth of the salinity maxima, and the depth of dissolved oxygen minima. Stations are organized from coastal to offshore (left to right).

**Figure S9**. Daily averages for discharge from the Mississippi River (A) across a ten year period and (B) during 2023. Dotted blue lines in (B) denote the duration of this study in 2023. Units for discharge are in feet^3^ second^-1^ (y-axes).

**Figure S10**. (A) Oil rig structures (symbols) in the Northern Gulf, where the green line with red circles indicates the route of our cruise survey. (B) withinX days of our survey, there was a detected oil spill near station 9. Grey line in (B) indicates the extent of the oil spill, based on satellite.

**Figure S11**. (A) Map of surface stations labeled with kmeans clusters determined using results from 18S rRNA gene metabarcoding.(B) Depth profile (y-axis) of all samples collected for this survey, mapped along all transects (y-axes), and labeled as kmeans clustering group.

**Figures S12-S15**. ASV richness by location (spatial scale) for individual taxonomic groups.

**Figure S16**. Total number of ASVs across ciliates and dinoflagellates (non-syndiniales and syndiniales only) assigned trophic modes (Jones et al. 2025). Bar plots report the specific feeding mode (morphotype) across depth (y-axis).

## References

Anderson, S. R., L. Blanco-Bercial, C. A. Carlson, and E. L. Harvey. 2024. Role of Syndiniales parasites in depth-specific networks and carbon flux in the oligotrophic ocean. ISME Commun. 4: ycae014.

Anglès, S., A. Jordi, D. W. Henrichs, and L. Campbell. 2019. Influence of coastal upwelling and river discharge on the phytoplankton community composition in the northwestern Gulf of Mexico. Prog. Oceanogr. 173: 26–36.

Application Note 31 Calculate Temperature and Conductivity Slope and Offset Correction Coefficients. 2024.Sea-Bird Scientific.

Application Note 64-2 SBE 43 DO Sensor Calibration and Data Corrections. 2024.Sea-Bird Scientific.

Bachy, C., E. Hehenberger, Y.-C. Ling, D. M. Needham, J. Strauss, S. Wilken, and A. Z. Worden. 2022. Marine protists: A hitchhiker’s guide to their role in the marine microbiome, p. 159–241. In The Microbiomes of Humans, Animals, Plants, and the Environment. Springer International Publishing.

Beatty, J. L., B. P. Stewart, L. Y. Mesrop, E. F. DeLong, D. M. Karl, and D. A. Caron. 2025. Eddy dipole differentially influences particle-associated and water column protistan community composition. Limnol. Oceanogr. 70: 817–832.

Becker, S., M. Aoyama, E. M. S. Woodward, K. Bakker, S. Coverly, C. Mahaffey, and T. Tanhua. 2020. GO-SHIP repeat hydrography nutrient manual: The precise and accurate determination of dissolved inorganic nutrients in seawater, using continuous flow analysis methods. Front. Mar. Sci. 7. doi:10.3389/fmars.2020.581790

Berlinches de Gea, A., J. Walochnik, J. Boenigk, K. Dumack, F. Henriquez, S. Rückert, M. Simon, and S. Geisen. 2025. Protists as determinants of the One Health framework. ISME J. 19. doi:10.1093/ismejo/wraf179

Bi, R., Z. Cao, S. M. H. Ismar-Rebitz, U. Sommer, H. Zhang, Y. Ding, and M. Zhao. 2021. Responses of marine diatom-dinoflagellate competition to multiple environmental drivers: Abundance, elemental, and biochemical aspects. Front. Microbiol. 12: 731786.

Biard, T. 2022. Diversity and ecology of Radiolaria in modern oceans. Environ. Microbiol. 24: 2179–2200.

Biggs, D. C. 1992. Nutrients, plankton, and productivity in a warm-core ring in the western Gulf of Mexico. J. Geophys. Res. 97: 2143.

Boeuf, D. and others. 2019. Biological composition and microbial dynamics of sinking particulate organic matter at abyssal depths in the oligotrophic open ocean. Proceedings of the National Academy of Sciences 201903080.

Bolyen, E. and others. 2019. Reproducible, interactive, scalable and extensible microbiome data science using QIIME 2. Nat. Biotechnol. 37: 852–857.

Cai, W.-J. and others. 2011. Acidification of subsurface coastal waters enhanced by eutrophication. Nat. Geosci. 4: 766–770.

Callahan, B. J., P. J. McMurdie, M. J. Rosen, A. W. Han, A. J. A. Johnson, and S. P. Holmes. 2016. DADA2: High-resolution sample inference from Illumina amplicon data. Nat. Methods 13: 581–583.

Campbell, L. G., J. C. Thrash, N. N. Rabalais, and O. U. Mason. 2019. Extent of the annual Gulf of Mexico hypoxic zone influences microbial community structure. PLoS One 14: e0209055.

Caron, D. A., P. D. Countway, A. C. Jones, D. Y. Kim, and A. Schnetzer. 2012. Marine Protistan Diversity. Ann. Rev. Mar. Sci. 4: 467–493.

Chakraborty, S., and S. E. Lohrenz. 2015. Phytoplankton community structure in the river–influenced continental margin of the northern Gulf of Mexico. Mar. Ecol. Prog. Ser. 521: 31–47.

Chakraborty, S., S. E. Lohrenz, and K. Gundersen. 2017. Photophysiological and light absorption properties of phytoplankton communities in the river-dominated margin of the northern Gulf of Mexico. J. Geophys. Res. C: Oceans 122: 4922–4938.

Chambouvet, A., P. Morin, D. Marie, and L. Guillou. 2008. Control of toxic marine dinoflagellate blooms by serial parasitic killers. Science 322: 1254–1257.

Chang, Y.-L., and L.-Y. Oey. 2012. Why does the Loop Current tend to shed more eddies in summer and winter? Geophys. Res. Lett. 39. doi:10.1029/2011gl050773

Dagg, M. J., J. W. Ammerman, R. M. W. Amon, W. S. Gardner, R. E. Green, and S. E. Lohrenz. 2007. A review of water column processes influencing hypoxia in the northern Gulf of Mexico. Estuaries Coasts 30: 735–752.

Damien, P., J. Sheinbaum, O. Pasqueron de Fommervault, J. Jouanno, L. Linacre, and O. Duteil. 2021. Do Loop Current eddies stimulate productivity in the Gulf of Mexico? Biogeosciences 18: 4281–4303.

Davis, N. M., D. M. Proctor, S. P. Holmes, D. A. Relman, and B. J. Callahan. 2018. Simple statistical identification and removal of contaminant sequences in marker-gene and metagenomics data. Microbiome 6: 226.

E Selph, K. R. Swalethorp, M. R Stukel, T. B Kelly, A. N Knapp, K. Fleming, T. Hernandez, and M. R Landry. 2022. Phytoplankton community composition and biomass in the oligotrophic Gulf of Mexico. J. Plankton Res. 44: 618–637.

Edwards, K. F., M. K. Thomas, C. A. Klausmeier, and E. Litchman. 2015. Light and growth in marine phytoplankton: allometric, taxonomic, and environmental variation: Light and growth in marine phytoplankton. Limnol. Oceanogr. 60: 540–552.

Fawcett, S. E., and B. B. Ward. 2011. Phytoplankton succession and nitrogen utilization during the development of an upwelling bloom. Mar. Ecol. Prog. Ser. 428: 13–31.

Flores, E., D. K. Romanovicz, M. Nieves-Morión, R. A. Foster, and T. A. Villareal. 2022. Adaptation to an intracellular lifestyle by a nitrogen-fixing, heterocyst-forming cyanobacterial Endosymbiont of a diatom. Front. Microbiol. 13: 799362.

Giner, C. R., M. C. Pernice, V. Balagué, C. M. Duarte, J. M. Gasol, R. Logares, and R. Massana. 2019. Marked changes in diversity and relative activity of picoeukaryotes with depth in the world ocean. ISME J. doi:10.1038/s41396-019-0506-9

Gleich, S. J., S. K. Hu, A. I. Krinos, and D. A. Caron. 2024. Protistan community composition and metabolism in the North Pacific Subtropical Gyre: Influences of mesoscale eddies and depth. Environ. Microbiol. 26: e16556.

Guillou, L. and others. 2012. The Protist Ribosomal Reference database (PR2): a catalog of unicellular eukaryote Small Sub-Unit rRNA sequences with curated taxonomy. Nucleic Acids Res. 41: D597–D604.

Harke, M. J., K. R. Frischkorn, G. M. M. Hennon, S. T. Haley, B. Barone, D. M. Karl, and S. T. Dyhrman. 2021. Microbial community transcriptional patterns vary in response to mesoscale forcing in the North Pacific Subtropical Gyre. Environ. Microbiol. 23: 4807–4822.

Hartigan, J. A., and M. A. Wong. 1979. Algorithm AS 136: A K-means clustering algorithm. J. R. Stat. Soc. Ser. C. Appl. Stat. 28: 100.

Henson, M. W., and J. C. Thrash. 2024. Microbial ecology of northern Gulf of Mexico estuarine waters. mSystems 9: e0131823.

Herzka, S. Z. S. by O., G. Samperio-Ramos, O. Hernández-Sánchez, V. F. Camacho Ibar, J. M. S. by O. Martin Hernandez-Ayon, E. S. by O. Pallas Sanz, M. C. Tenreiro, and J. Sheinbaum. 2024. Contribution of subsurface nitrate and fixed nitrogen inside and outside of a Loop Current anticyclonic eddy based on zooplankton stable isotope ratios. Proceedings of the Ocean Sciences Meeting.

Hood, E. M., C. L. Sabine, and B. M. Sloyan. 2010. The GO-SHIP Repeat Hydrography Manual: A Collection of Expert Reports and Guidelines. IOCCP Report 14. IOCCP Report 14.

Hu, S. K., Paige E. Connell, L. Y. Mesrop, and D. A. Caron. 2018. A Hard Day’s Night: Diel Shifts in Microbial Eukaryotic Activity in the North Pacific Subtropical Gyre. Frontiers in Marine Science 5: 351.

Jephcott, T. G., C. Alves-de-Souza, F. H. Gleason, F. F. van Ogtrop, T. Sime-Ngando, S. A. Karpov, and L. Guillou. 2016. Ecological impacts of parasitic chytrids, syndiniales and perkinsids on populations of marine photosynthetic dinoflagellates. Fungal Ecol. 19: 47–58.

Johansson, M. 2004. Annual variability in ciliate community structure, potential prey and predators in the open northern Baltic Sea proper. J. Plankton Res. 26: 67–80.

Jones, E. L., S. Menden-Deuer, and T. Rynearson. 2025. Trophic Mode Database (TMD) for Dinoflagellate and Ciliate Species.doi:10.5281/ZENODO.15149504

Jones-Kellett, A. E., J. C. McNichol, Y. Raut, J. A. Fuhrman, and M. J. Follows. 2026. The dynamic mesoscale sink and source niches for eukaryotic phytoplankton in a subtropical gyre. Proc. Natl. Acad. Sci. U. S. A. 123: e2608700123.

Karl, D. M., M. J. Church, J. E. Dore, R. M. Letelier, and C. Mahaffey. 2012. Predictable and efficient carbon sequestration in the North Pacific Ocean supported by symbiotic nitrogen fixation. Proceedings of the National Academy of Sciences 109: 1842–1849.

Kemp, A. E. S., and T. A. Villareal. 2018. The case of the diatoms and the muddled mandalas: Time to recognize diatom adaptations to stratified waters. Prog. Oceanogr. 167: 138–149.

Krause, J. W., A. D. Boyette, I. A. Marquez, R. A. Pickering, and K. Maiti. 2023. Drivers of diatom production and the legacy of eutrophication in two river plume regions of the northern Gulf of Mexico. Front. Mar. Sci. 10: 1162685.

Landry, M. R., K. E. Selph, M. R. Stukel, R. Swalethorp, T. B. Kelly, J. L. Beatty, and C. R. Quackenbush. 2022. Microbial food web dynamics in the oceanic Gulf of Mexico. J. Plankton Res. 44: 638–655.

Langdon, C. 2010. Determination of dissolved oxygen in seawater by Winkler titration using the amperometric technique.

Lee, S., G. Wolberg, and S. Y. Shin. 1997. Scattered data interpolation with multilevel B-splines. IEEE Trans. Vis. Comput. Graph. 3: 228–244.

Lima-Mendez, G. and others. 2015. Determinants of community structure in the global plankton interactome. Science 348: 1262073–1262073.

Linacre, L. and others. 2025. Carbon biomass assessment for microbial plankton groups across a young anticyclonic loop current eddy and surrounding waters in the Gulf of Mexico. J. Geophys. Res. Oceans 130: e2025JC022705.

Liu, Y., Z. Hu, Y. Deng, L. Shang, C. J. Gobler, and Y. Z. Tang. 2021. Dependence of genome size and copy number of rRNA gene on cell volume in dinoflagellates. Harmful Algae 109: 102108.

Lohrenz, S. E., D. G. Redalje, W.-J. Cai, J. Acker, and M. Dagg. 2008. A retrospective analysis of nutrients and phytoplankton productivity in the Mississippi River plume. Cont. Shelf Res. 28: 1466–1475.

Maiti, K., S. Bosu, E. J. D’Sa, P. L. Adhikari, M. Sutor, and K. Longnecker. 2016. Export fluxes in northern Gulf of Mexico - Comparative evaluation of direct, indirect and satellite-based estimates. Mar. Chem. 184: 60–77.

Mars Brisbin, M., O. D. Brunner, M. M. Grossmann, and S. Mitarai. 2020. Paired high-throughput, in situ imaging and high-throughput sequencing illuminate acantharian abundance and vertical distribution. Limnol. Oceanogr. 65: 2953–2965.

Martin, M. 2011. Cutadapt removes adapter sequences from high-throughput sequencing reads. EMBnet.journal 17: 10–12.

Mason, O. U., E. J. Canter, L. E. Gillies, T. K. Paisie, and B. J. Roberts. 2016. Mississippi River plume enriches microbial diversity in the northern Gulf of Mexico. Front. Microbiol. 7: 1048.

McGillicuddy, D. J., Jr and others. 2007. Eddy/wind interactions stimulate extraordinary mid-ocean plankton blooms. Science 316: 1021–1026.

McGillicuddy, D. J., Jr. 2016. Mechanisms of physical-biological-biogeochemical interaction at the oceanic mesoscale. Ann. Rev. Mar. Sci. 8: 125–159.

Millette, N. C. and others. 2023. Mixoplankton and mixotrophy: future research priorities. 45: 576–596.

Mitra, A. and others. 2023. The Mixoplankton Database (MDB): Diversity of photo-phago-trophic plankton in form, function, and distribution across the global ocean. J. Eukaryot. Microbiol. e12972.

Officer, C. B., and J. H. Ryther. 1980. The possible importance of silicon in marine eutrophication. Mar. Ecol. Prog. Ser. 3: 83–91.

Oksanen, J., R. Kindt, P. Legendre, B. O’Hara, M. H. H. Stevens, M. J. Oksanen, and M. Suggests. 2007. The vegan package. Community ecology package 10: 719.

Ollison, G. A., S. K. Hu, L. Y. Mesrop, E. F. DeLong, and D. A. Caron. 2021. Come rain or shine: Depth not season shapes the active protistan community at station ALOHA in the North Pacific Subtropical Gyre. Deep Sea Res. Part 1 Oceanogr. Res. Pap. 170: 103494.

Payne, R. 2013. Seven reasons why protists make useful bioindicators. Acta Protozoologica 2013: 105–113.

Portela, E., M. Tenreiro, E. Pallàs-Sanz, T. Meunier, A. Ruiz-Angulo, R. Sosa-Gutiérrez, and S. Cusí. 2018. Hydrography of the central and western gulf of Mexico. J. Geophys. Res. Oceans 123: 5134–5149.

Pyle, A. E., A. M. Johnson, and T. A. Villareal. 2020. Isolation, growth, and nitrogen fixation rates of the Hemiaulus-Richelia (diatom-cyanobacterium) symbiosis in culture. PeerJ 8: e10115.

Rabalais, N. N., R. E. Turner, and W. J. Wiseman Jr. 2002. Gulf of Mexico hypoxia, A.k.a. “the dead zone.” Annu. Rev. Ecol. Syst. 33: 235–263.

Rii, Y., D. Karl, and M. Church. 2016. Temporal and vertical variability in picophytoplankton primary productivity in the North Pacific Subtropical Gyre. Mar. Ecol. Prog. Ser. 562: 1–18.

Rognes, T., T. Flouri, B. Nichols, C. Quince, and F. Mahé. 2016. VSEARCH: a versatile open source tool for metagenomics. PeerJ 4: e2584.

Sandin, M. M., J. Renaudie, N. Suzuki, and F. Not. 2025. Extant diversity, biogeography, and evolutionary history of Radiolaria. Curr. Biol. 35: 2524–2538.e6.

Santoferrara, L. F., A. Qureshi, A. Sher, and L. Blanco-Bercial. 2023. The photic-aphotic divide is a strong ecological and evolutionary force determining the distribution of ciliates (Alveolata, Ciliophora) in the ocean. J. Eukaryot. Microbiol. 70: e12976.

Sidón-Ceseña, K., M. A. Martínez-Mercado, J. Chong-Robles, Y. Ortega-Saad, V. F. Camacho-Ibar, L. Linacre, and A. Lago-Lestón. 2025. The protist community of the oligotrophic waters of the Gulf of Mexico is distinctly shaped by depth-specific physicochemical conditions during the warm season. FEMS Microbiol. Ecol. 101: fiaf009.

Snyder, R. A., J. A. Moss, L. Santoferrara, M. Head, and W. Jeffrey. 2021. Ciliate microzooplankton from the northeastern Gulf of Mexico. Ices Journal of Marine Science 78: 3356–3371.

Stoeck, T., D. Bass, M. Nebel, R. Christen, M. D. M. Jones, H.-W. Breiner, and T. A. Richards. 2010. Multiple marker parallel tag environmental DNA sequencing reveals a highly complex eukaryotic community in marine anoxic water. Mol. Ecol. 19: 21–31.

Sturges, W., and J. C. Evans. 1983. On the variability of the Loop Current in the Gulf of Mexico. J. Mar. Res. 41: 639–653.

Sturm, D., J. de Vries, W. M. Balch, G. Wheeler, and C. Brownlee. 2023. Mesoscale oceanographic meanders influence protist community function and structure in the southern Indian Ocean. Environ. Microbiol. 25: 3161–3179.

Tréguer, P. and others. 2018. Influence of diatom diversity on the ocean biological carbon pump. Nat. Geosci. 11: 27–37.

Turner, R. E., N. Qureshi, N. N. Rabalais, Q. Dortch, D. Justić, R. F. Shaw, and J. Cope. 1998. Fluctuating silicate:nitrate ratios and coastal plankton food webs. Proc. Natl. Acad. Sci. U. S. A. 95: 13048–13051.

Vaulot, D. 2021. pr2database/pr2database: PR2 version 4.14.0,.

Wang, Z. A., R. Wanninkhof, W.-J. Cai, R. H. Byrne, X. Hu, T.-H. Peng, and W.-J. Huang. 2013. The marine inorganic carbon system along the Gulf of Mexico and Atlantic coasts of the United States: Insights from a transregional coastal carbon study. Limnol. Oceanogr. 58: 325–342.

Ward, B. A., and M. J. Follows. 2016. Marine mixotrophy increases trophic transfer efficiency, mean organism size, and vertical carbon flux. Proceedings of the National Academy of Sciences 113: 2958–2963.

Wawrik, B., and J. H. Paul. 2004. Phytoplankton community structure and productivity along the axis of the Mississippi River plume in oligotrophic Gulf of Mexico waters. Aquat. Microb. Ecol. 35: 185–196.

Wawrik, B., J. H. Paul, L. Campbell, D. Griffin, L. Houchin, A. Fuentes-Ortega, and F. Muller-Karger. 2003. Vertical structure of the phytoplankton community associated with a coastal plume in the Gulf of Mexico. Mar. Ecol. Prog. Ser. 251: 87–101.

Wickham, H. 2017. The tidyverse. R package ver 1: 1.

Williams, A. K., A. S. McInnes, J. R. Rooker, and A. Quigg. 2015. Changes in microbial plankton assemblages induced by mesoscale oceanographic features in the Northern Gulf of Mexico. PLoS One 10: e0138230.

Worden, A. Z., M. J. Follows, S. J. Giovannoni, S. Wilken, A. E. Zimmerman, and P. J. Keeling. 2015. Rethinking the marine carbon cycle: Factoring in the multifarious lifestyles of microbes. Science 347: 1257594–1257594.

Wu, W., L. Wang, Y. Liao, and B. Huang. 2015. Microbial eukaryotic diversity and distribution in a river plume and cyclonic eddy-influenced ecosystem in the South China Sea. Microbiologyopen 4: 826–840.

Anderson, S. R., L. Blanco-Bercial, C. A. Carlson, and E. L. Harvey. 2024. Role of Syndiniales parasites in depth-specific networks and carbon flux in the oligotrophic ocean. ISME Communications, 4(1):ycae014,.

Anglès, S., A. Jordi, D. W. Henrichs, and L. Campbell. 2019. Influence of coastal upwelling and river discharge on the phytoplankton community composition in the northwestern Gulf of Mexico. Progress in Oceanography, 173:26–36,.

1971. Meteorological and Oceanographic Data Collected from the National Data Buoy Center Coastal-Marine Automated Network (C-MAN) and Moored (Weather) Buoys. pp.

2025. National Water Information System Data for Site 07374000 (U.S. Geological Survey, ed). Daily discharge, Mississippi River at Baton Rouge, LA, 2021–2025 pp.

