## Supplementary Figures for "Riverine input and eddy edge effects on microeukaryotic biodiversity in the Northern Gulf"

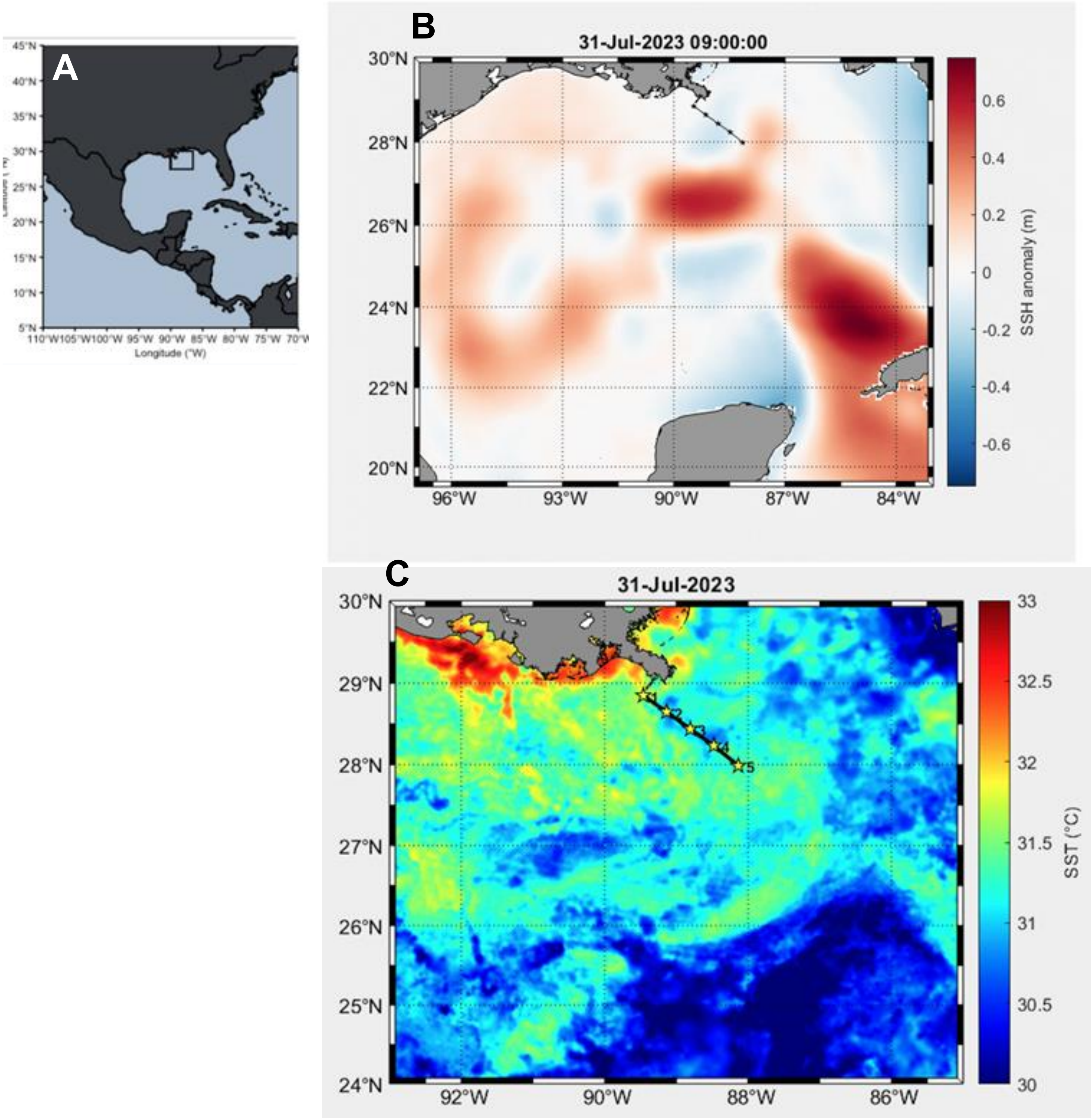


**Figure S1**. (**A**) Regional map showing the southeast United States and Mexico. The rectangle outlines the study area shown in B and C. (**B**) Sea surface height (SSH; m) of study area. Data points show the location of stations 1-5. (**C**) Sea surface temperature (°C) during the study period. Data points show the location of stations 1-5.

**Transect 1**

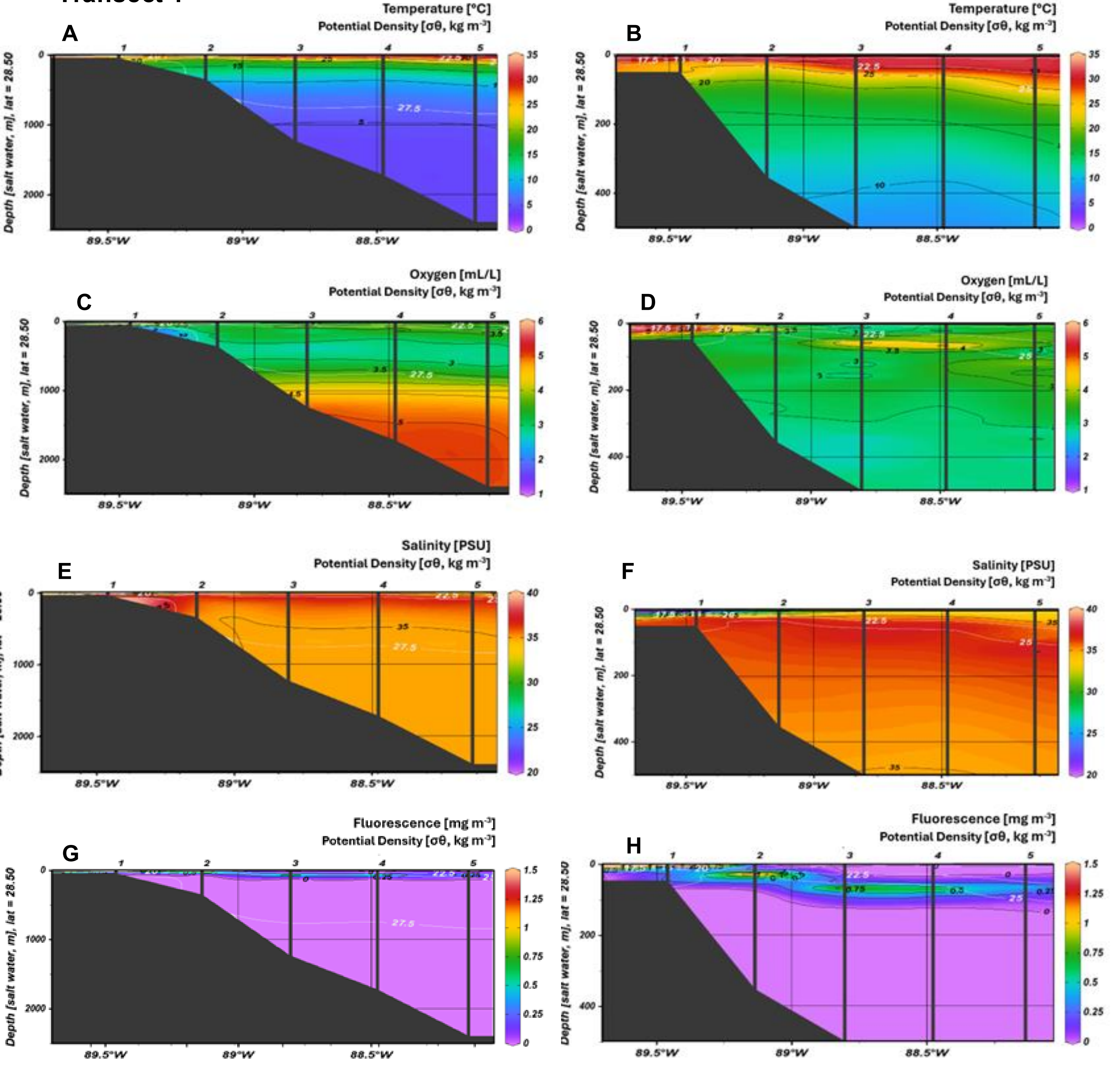


**Figure S2**. Oceanographic conditions at Transect 1. Stations are noted at the top of each panel with a vertical line through the profile and the potential density is mapped on top of each profile. (A) Temperature (°C) from the surface to deepest depth, and (B) from the surface to 500 m. (C) Oxygen (ml/L) from the surface to deepest depth and (D) from the surface to 500 m. (E) Salinity (PSU) for the full vertical profile and (F) up to 500 m. (G) Fluorescence (mg $m^{-3}$) throughout the full profile and (H) up to 500 m.

### Transect 2

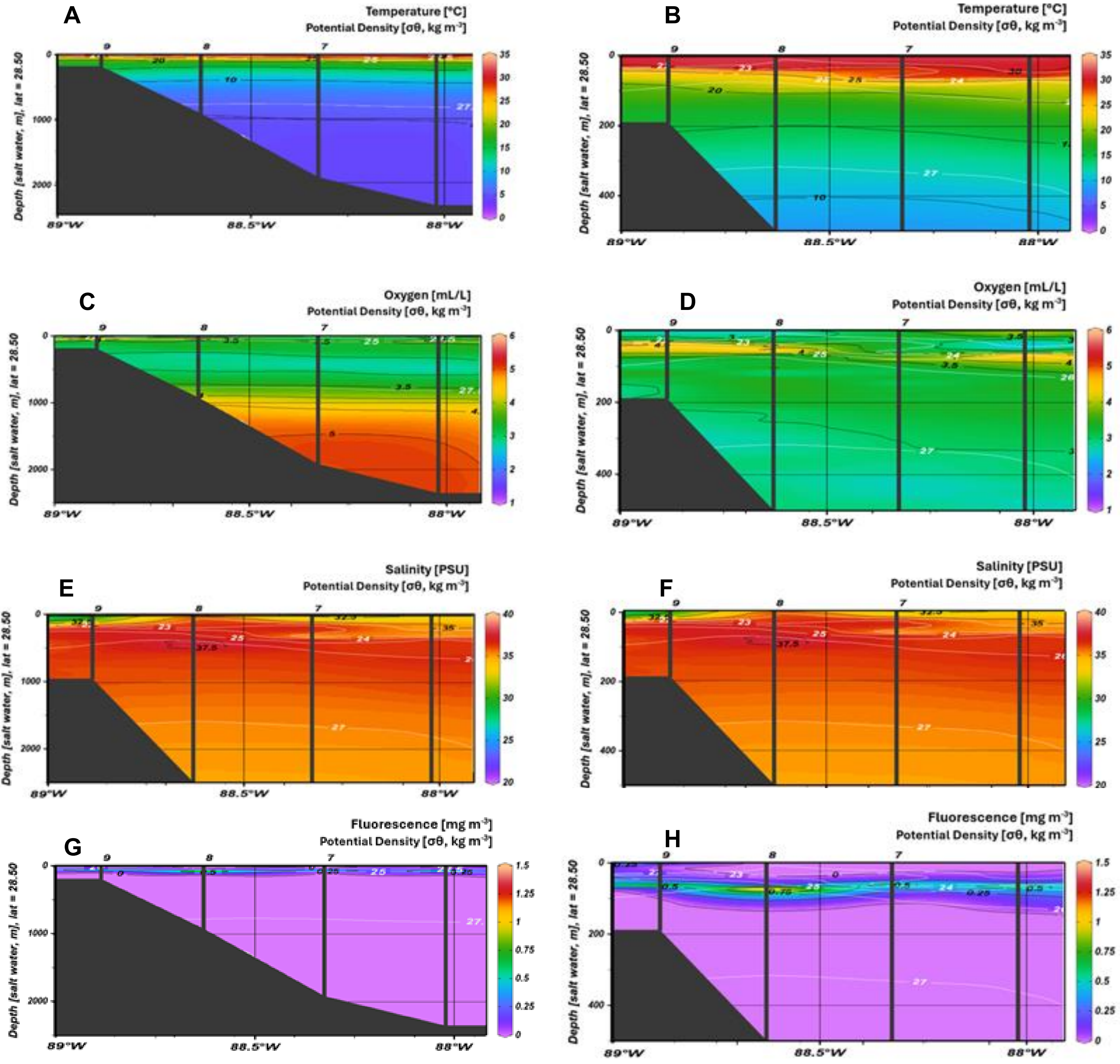


**Figure S3.** Vertical profiles along Transect 2. Stations are noted at the top of each panel with a vertical line through the profile and the potential density is mapped on top of each profile. (A) Temperature (°C) from the surface to deepest depth, and (B) from the surface to 500 m. (C) Oxygen (ml/L) from the surface to deepest depth and (D) from the surface to 500 m. (E) Salinity (PSU) for the full vertical profile and (F) up to 500 m. (G) Fluorescence (mg $m^{-3}$) throughout the full profile and (H) up to 500 m.

Transect 3

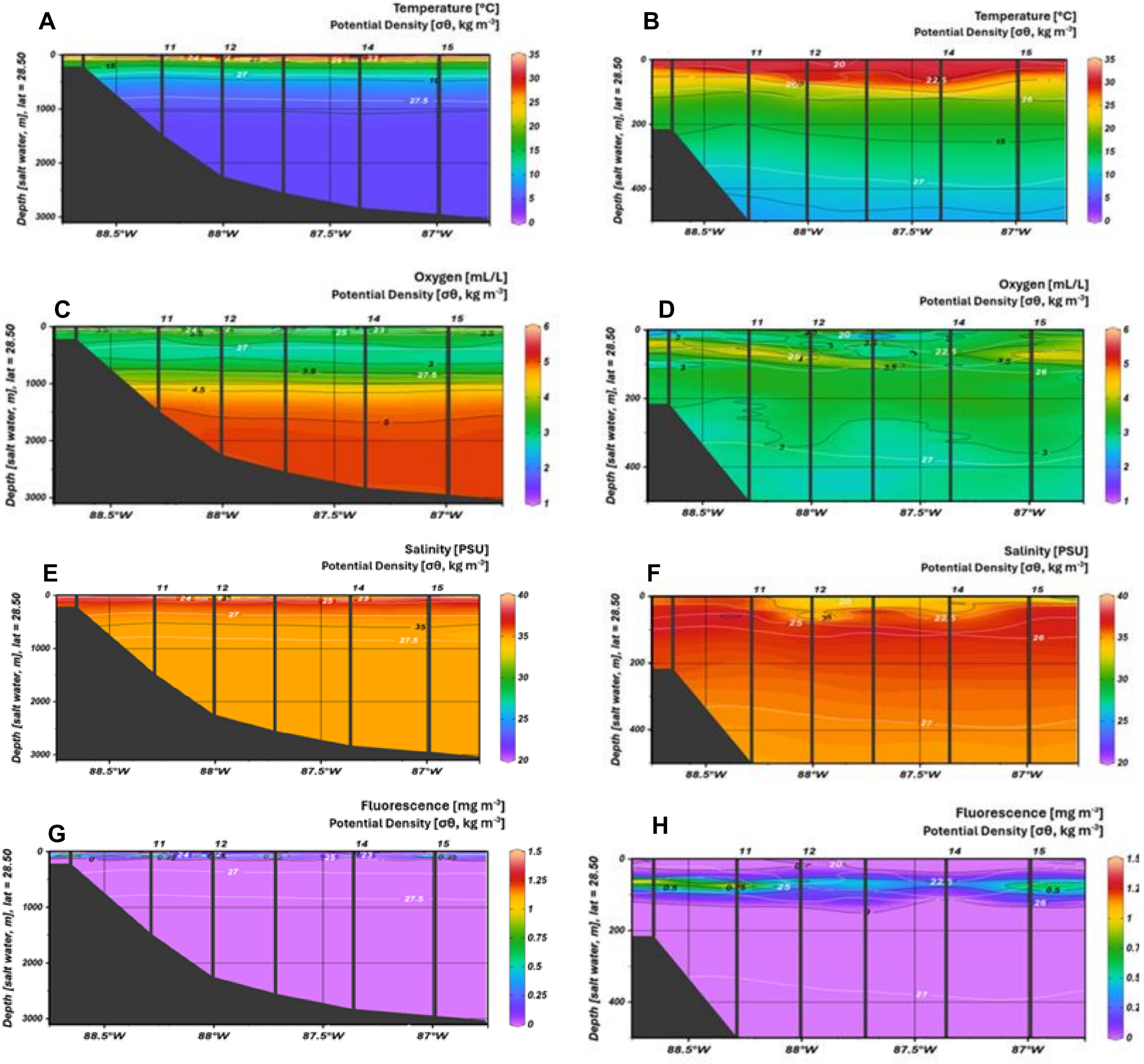


**Figure S4**. Transect 3 water column parameters. Stations are noted at the top of each panel with a vertical line through the profile and the potential density is mapped on top of each profile. (A) Temperature (°C) from the surface to deepest depth, and (B) from the surface to 500 m. (C) Oxygen (ml/L) from the surface to deepest depth and (D) from the surface to 500 m. (E) Salinity (PSU) for the full vertical profile and (F) up to 500 m. (G) Fluorescence (mg m$^{-3}$) throughout the full profile and (H) up to 500 m.

### Transect 1

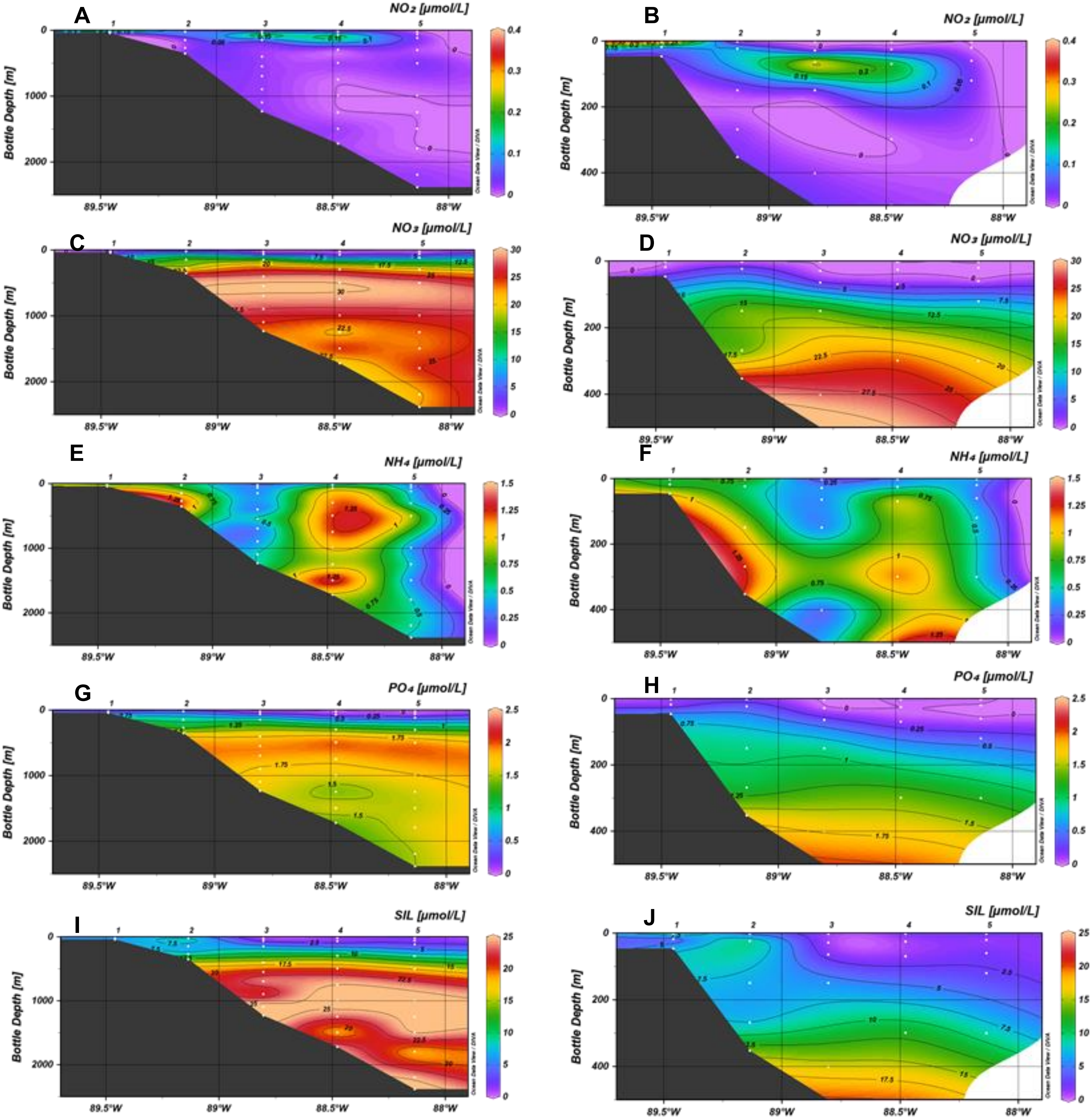


**Figure S5**. Nutrient profiles at Transect 1. ODV interpolation derived from discrete samples (white symbols) from each station (numeric at top of each panel). All units are in µmol $L^{-1}$ and the full profile (surface to deepest depth) is shown on the left-hand side panels and paired with the same profile up to 500 m on the right-hand side. Profiles are shown for (A-B) nitrite, $NO_2$, (C-D) nitrate, $NO_3$ , (E-F) ammonium, $NH_4$, (G-H) phosphate, $PO_4$ (I-J) and silicic acid (SIL).

### Transect 2

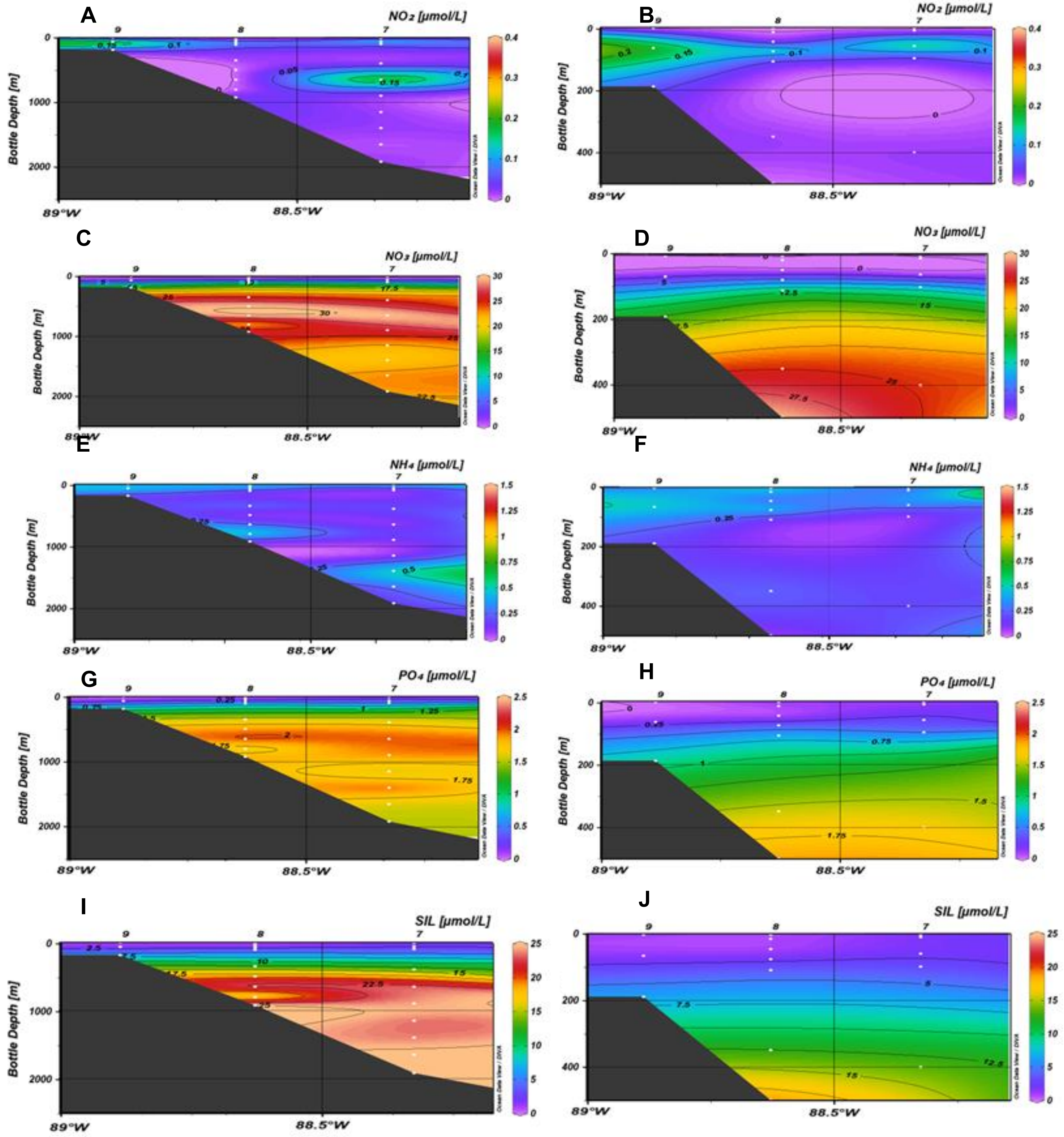


**Figure S6**. Nutrient profiles at Transect 2. ODV interpolation derived from discrete samples (white symbols) from each station (numeric at top of each panel). All units are in µmol $L^{-1}$ and the full profile (surface to deepest depth) is shown on the left-hand side panels and paired with the same profile up to 500 m on the right-hand side. Profiles are shown for (A-B) nitrite, $NO_2$, (C-D) nitrate, $NO_3$ , (E-F) ammonium, $NH_4$, (G-H) phosphate, $PO_4$ (I-J) and silicic acid (SIL).

### Transect 3

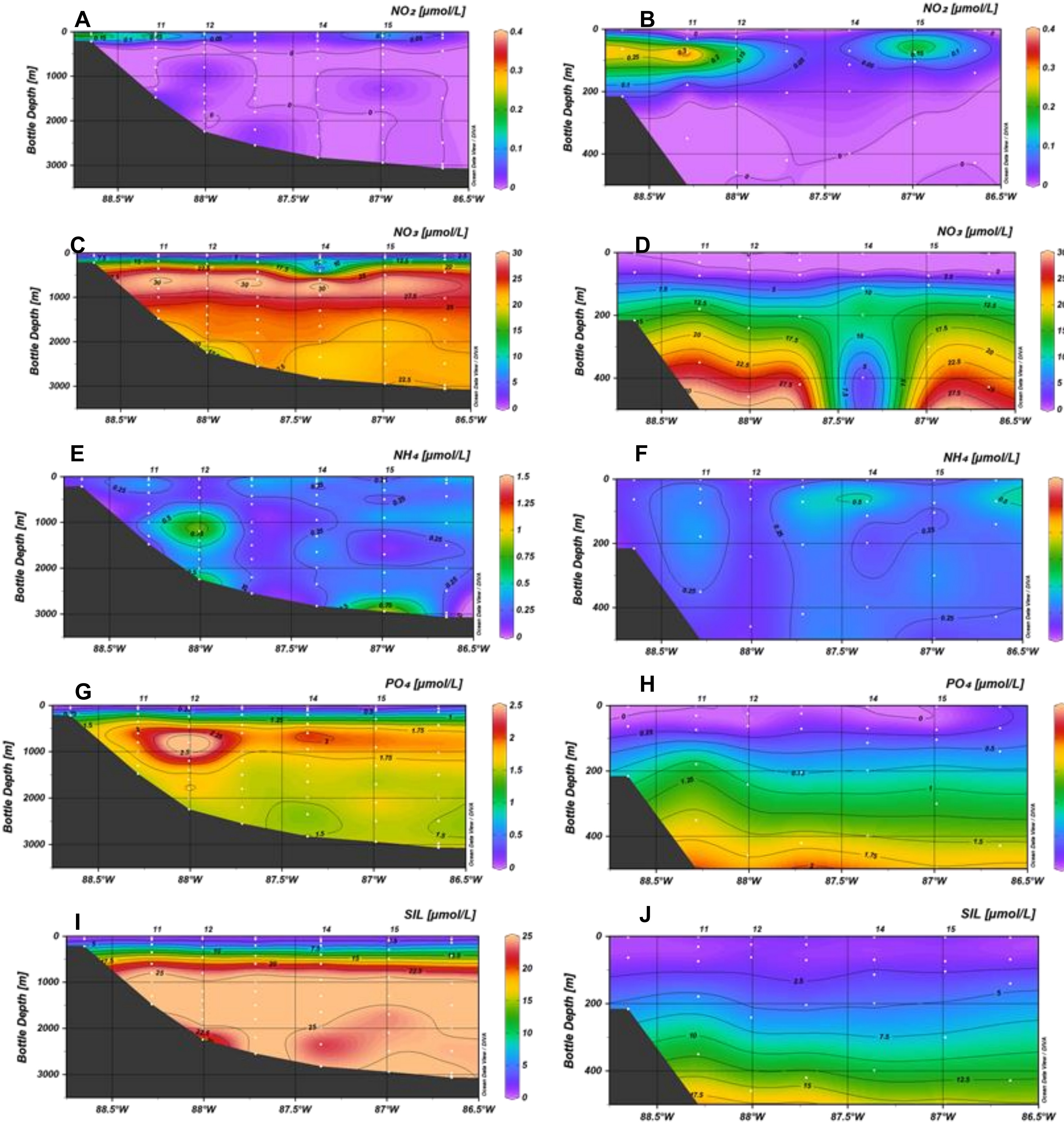


**Figure S7**. Nutrient profiles at Transect 3. ODV interpolation derived from discrete samples (white symbols) from each station (numeric at top of each panel). All units are in µmol $L^{-1}$ and the full profile (surface to deepest depth) is shown on the left-hand side panels and paired with the same profile up to 500 m on the right-hand side. Profiles are shown for (A-B) nitrite, $NO_2$, (C-D) nitrate, $NO_3$ , (E-F) ammonium, $NH_4$, (G-H) phosphate, $PO_4$ (I-J) and silicic acid (SIL).

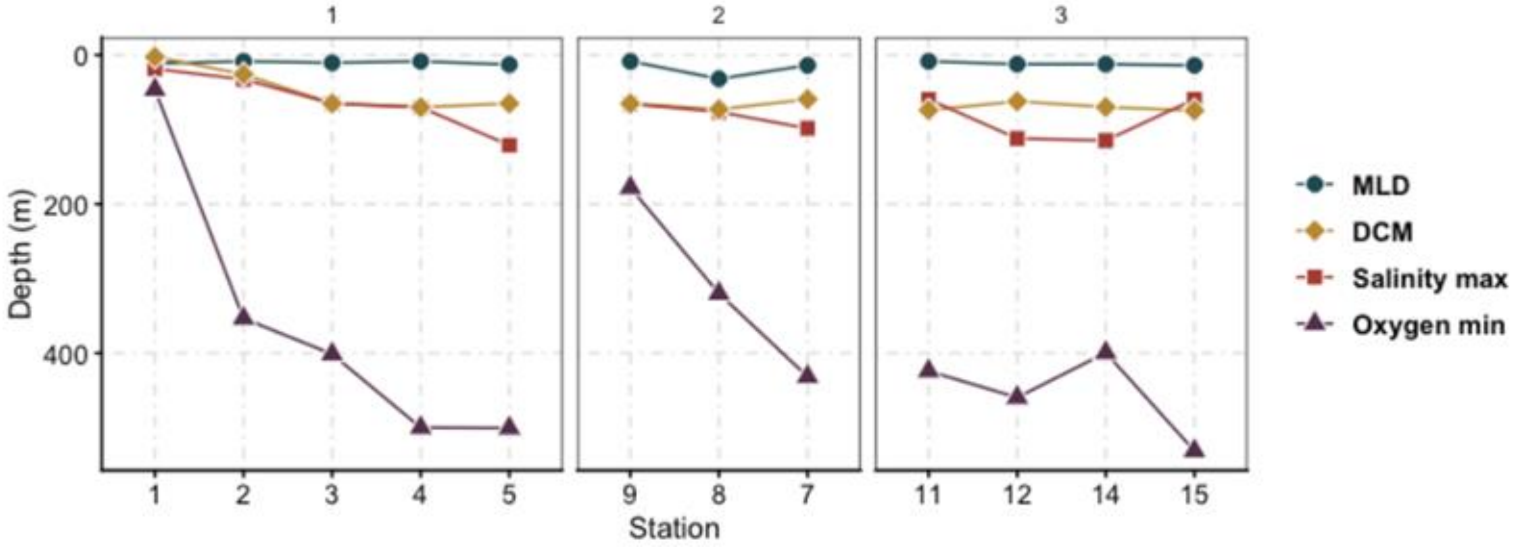


**Figure S8**. Depth features across each transect (panels from left to right) and station (x-axis bottom labels). Symbols and colors denote the mixed layer depth (MLD), deep chlorophyll maxima (DCM), depth of the salinity maxima, and the depth of dissolved oxygen minima. Stations are organized from coastal to offshore (left to right).

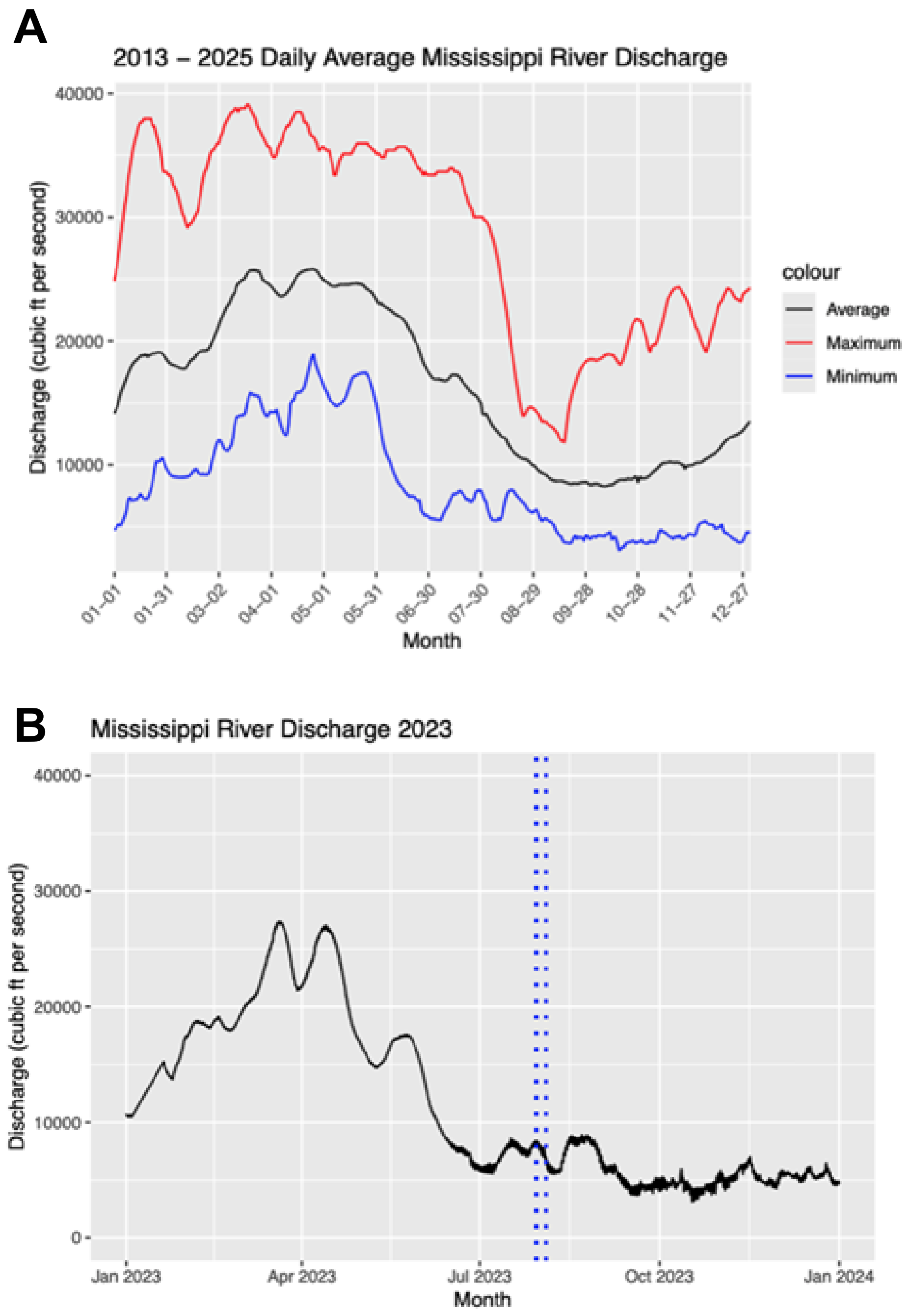


**Figure S9**. Daily averages for discharge from the Mississippi River (A) across a ten year period and (B) during 2023. Dotted blue lines in (B) denote the duration of this study in 2023. Units for discharge are in feet$^3$ second$^{-1}$ (y-axes).

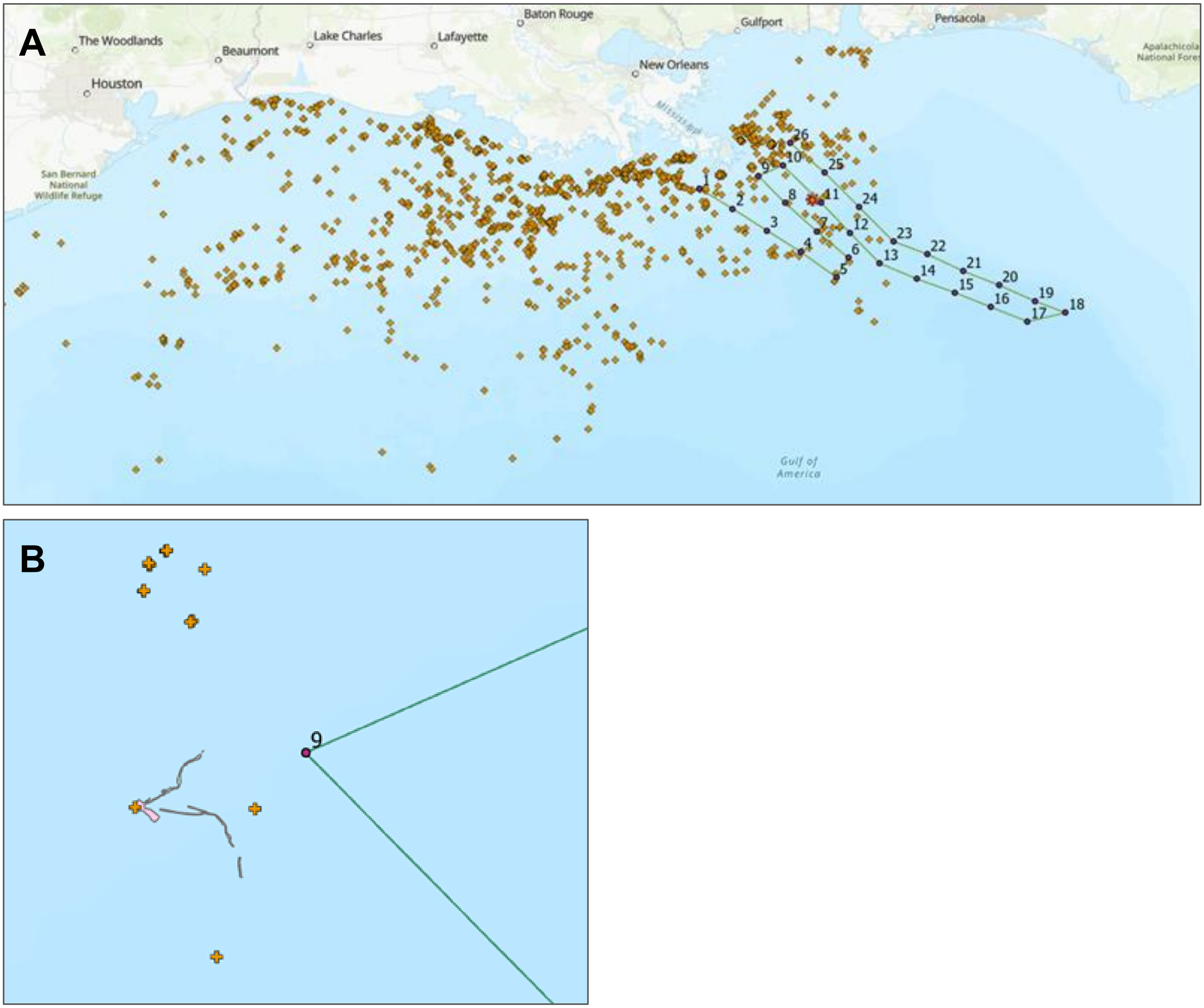


**Figure S10**. (A) Oil rig structures (symbols) in the Northern Gulf, where the green line with red circles indicates the route of our cruise survey. (B) withinX days of our survey, there was a detected oil spill near station 9. Grey line in (B) indicates the extent of the oil spill, based on satellite.

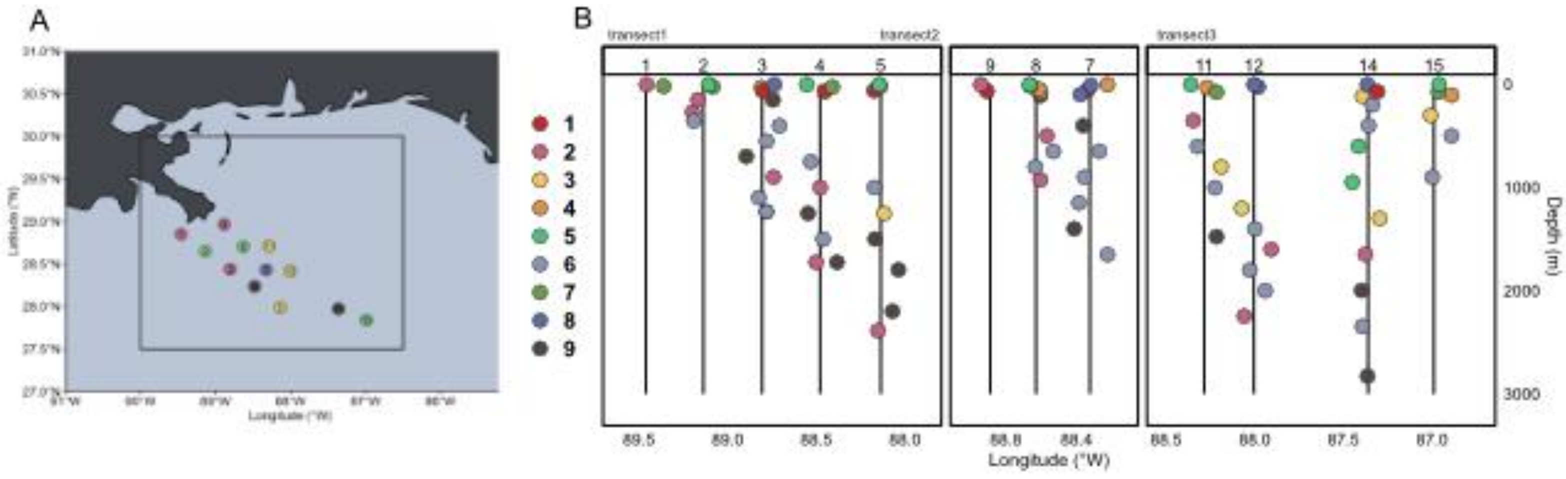


**Figure S11**. (A) Map of surface stations labeled with kmeans clusters determined using results from 18S rRNA gene metabarcoding. (B) Depth profile (y-axis) of all samples collected for this survey, mapped along all transects (y-axes), and labeled as kmeans clustering group.

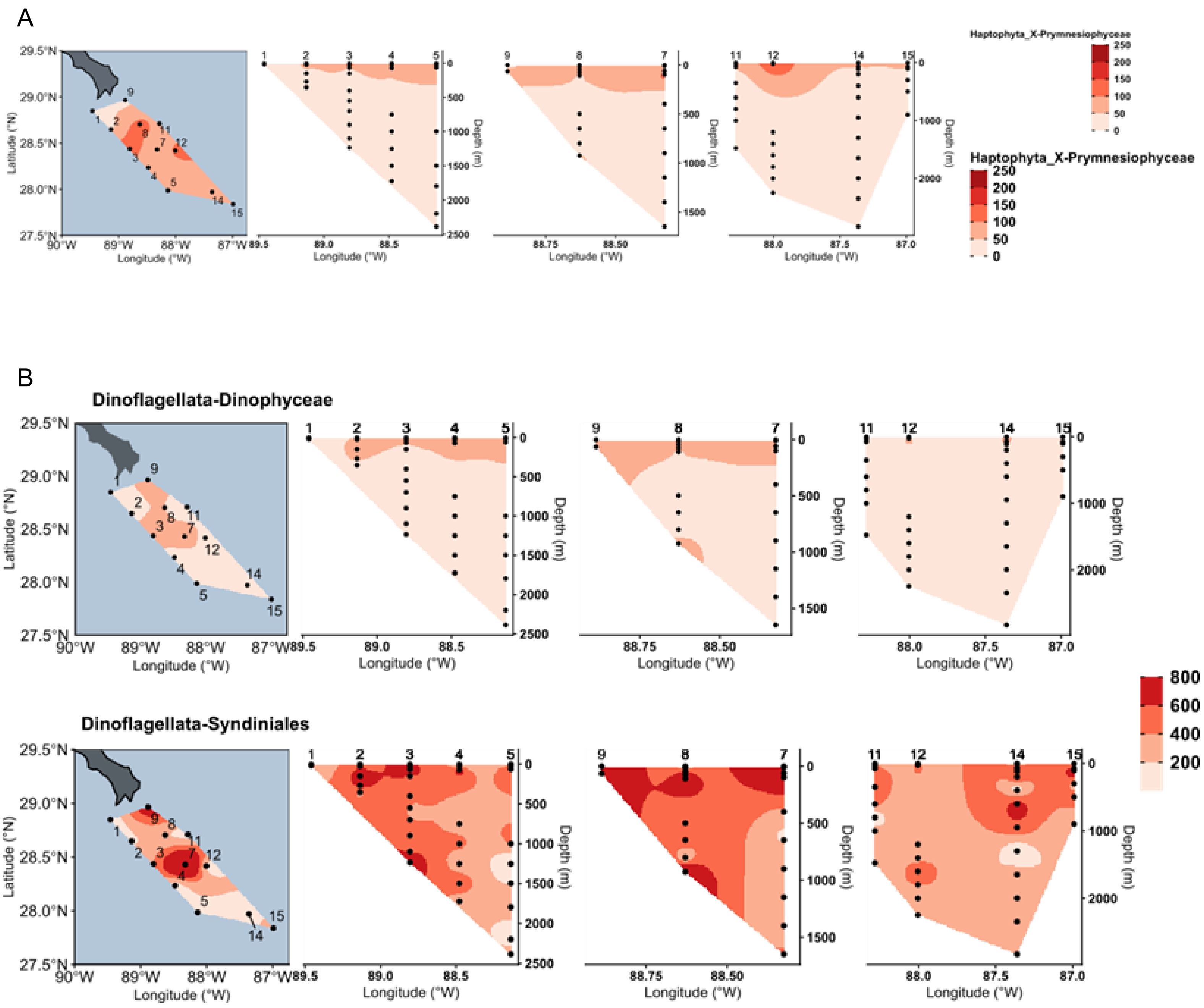


**Figure S12**. ASV richness for the (**A**) haptophyte group, Prymnesiophyceae and (**B**) dinophyceae (top) and Syndiniales (bottom) at the surface and throughout each transect. Mapping of ASVs spatially was interpolated by determining using a multilevel B-spline between points.

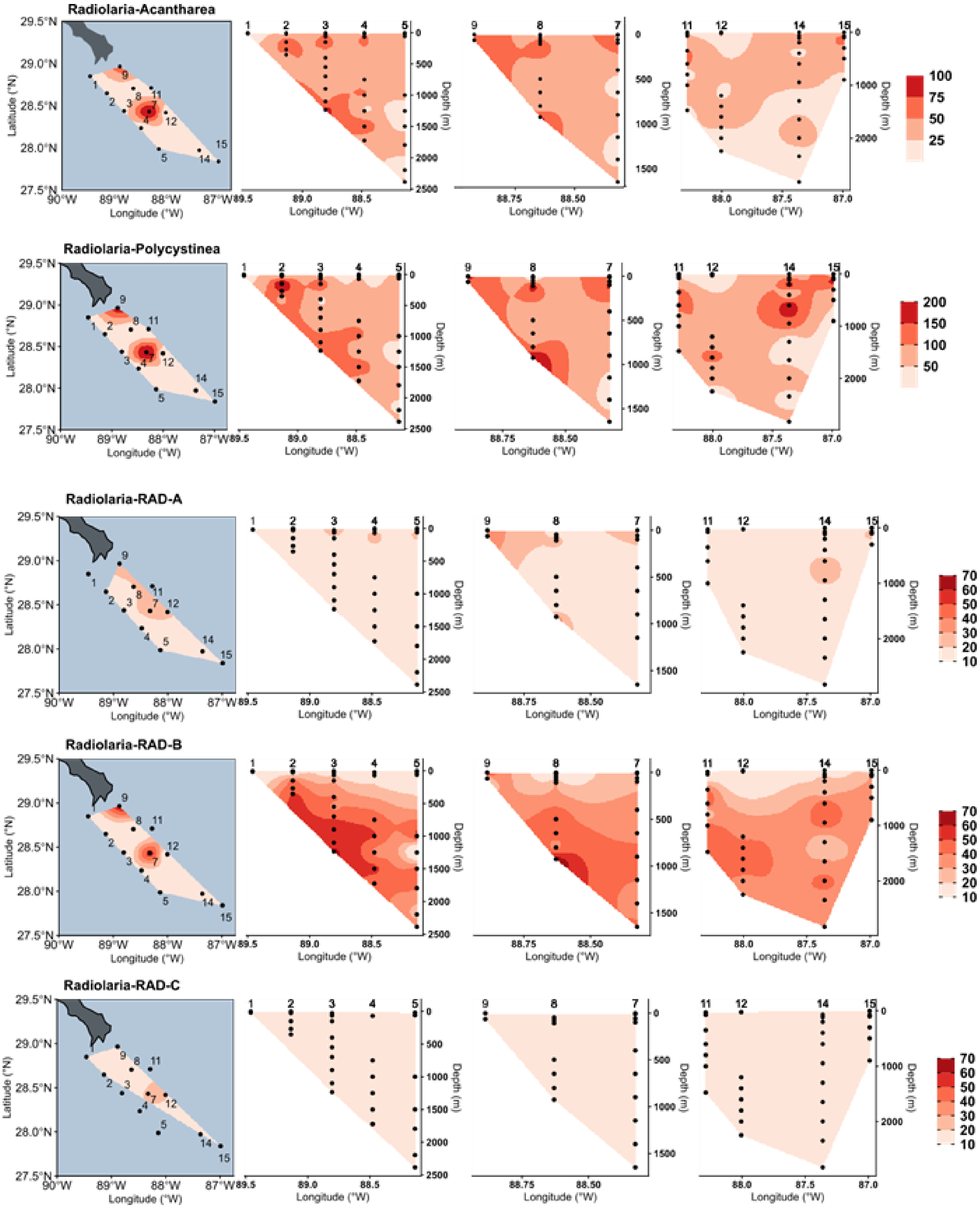


**Figure S13**. ASV richness for radiolarian groups at the surface and throughout each transect. Mapping of ASVs spatially was interpolated by determining using a multilevel B-spline between points.

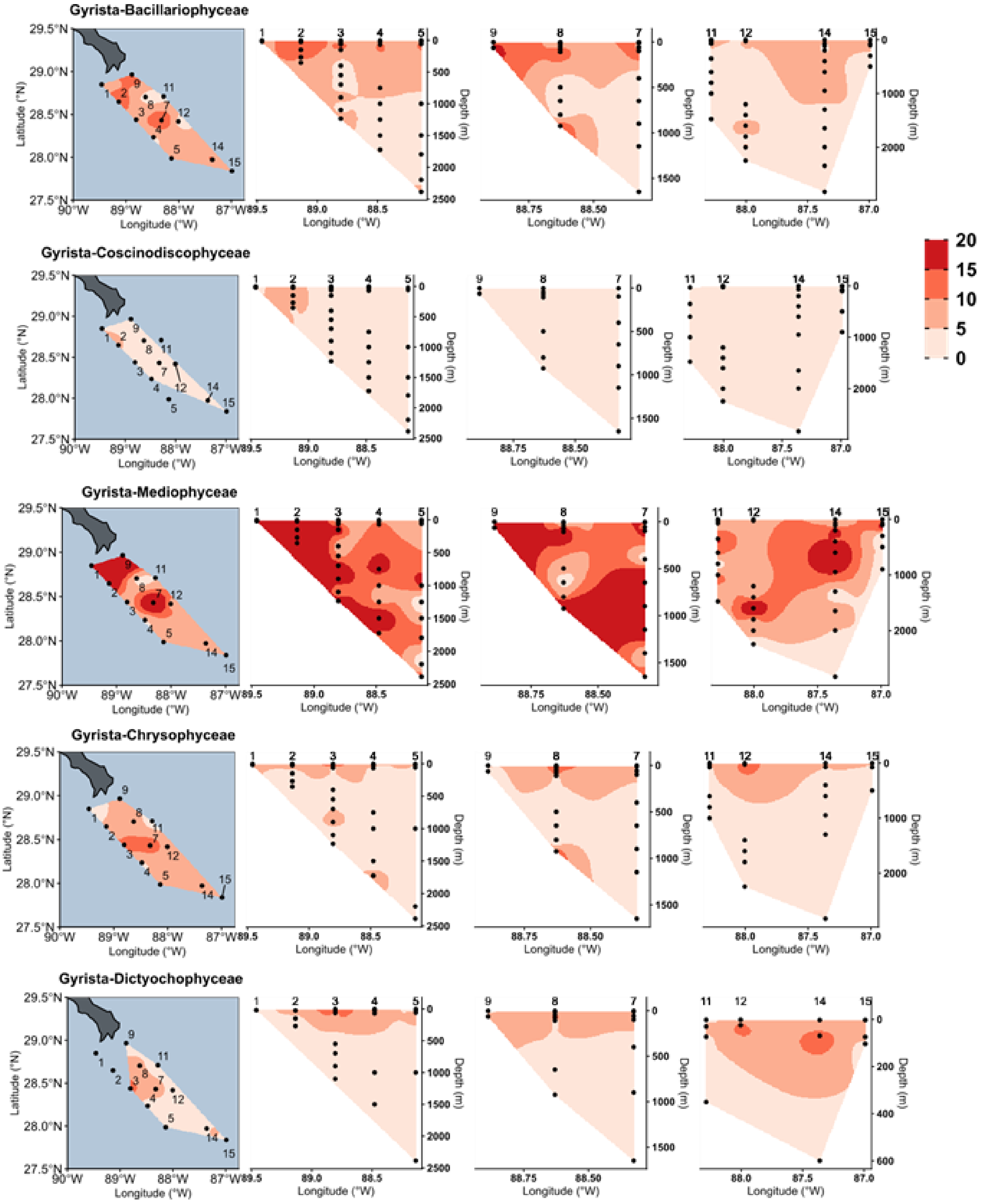


**Figure S14**. ASV richness for gyrista groups at the surface and throughout each transect. Mapping of ASVs spatially was interpolated by determining using a multilevel B-spline between points.

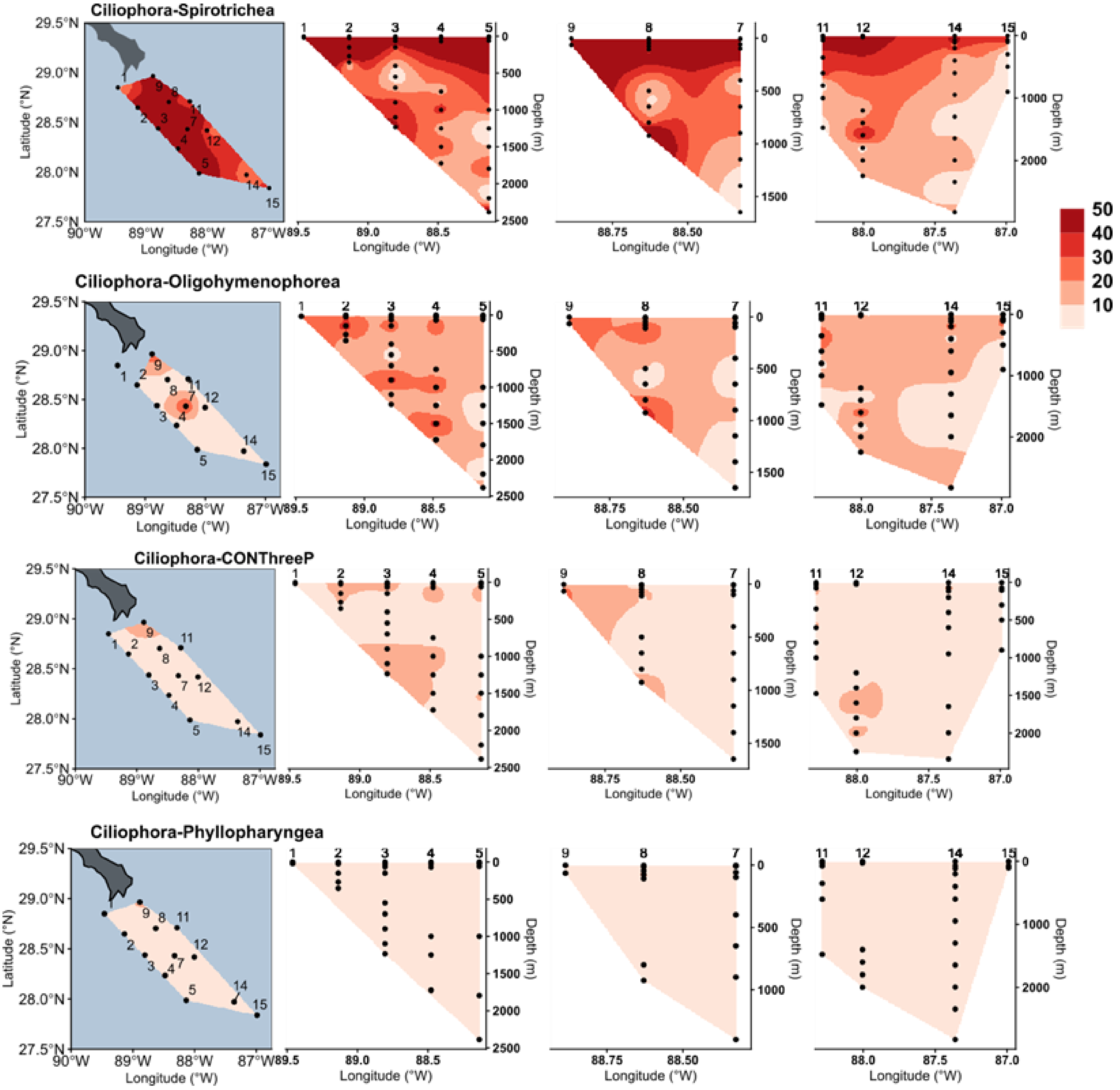


**Figure S15**. ASV richness for ciliate groups at the surface and throughout each transect. Mapping of ASVs spatially was interpolated by determining using a multilevel B-spline between points.

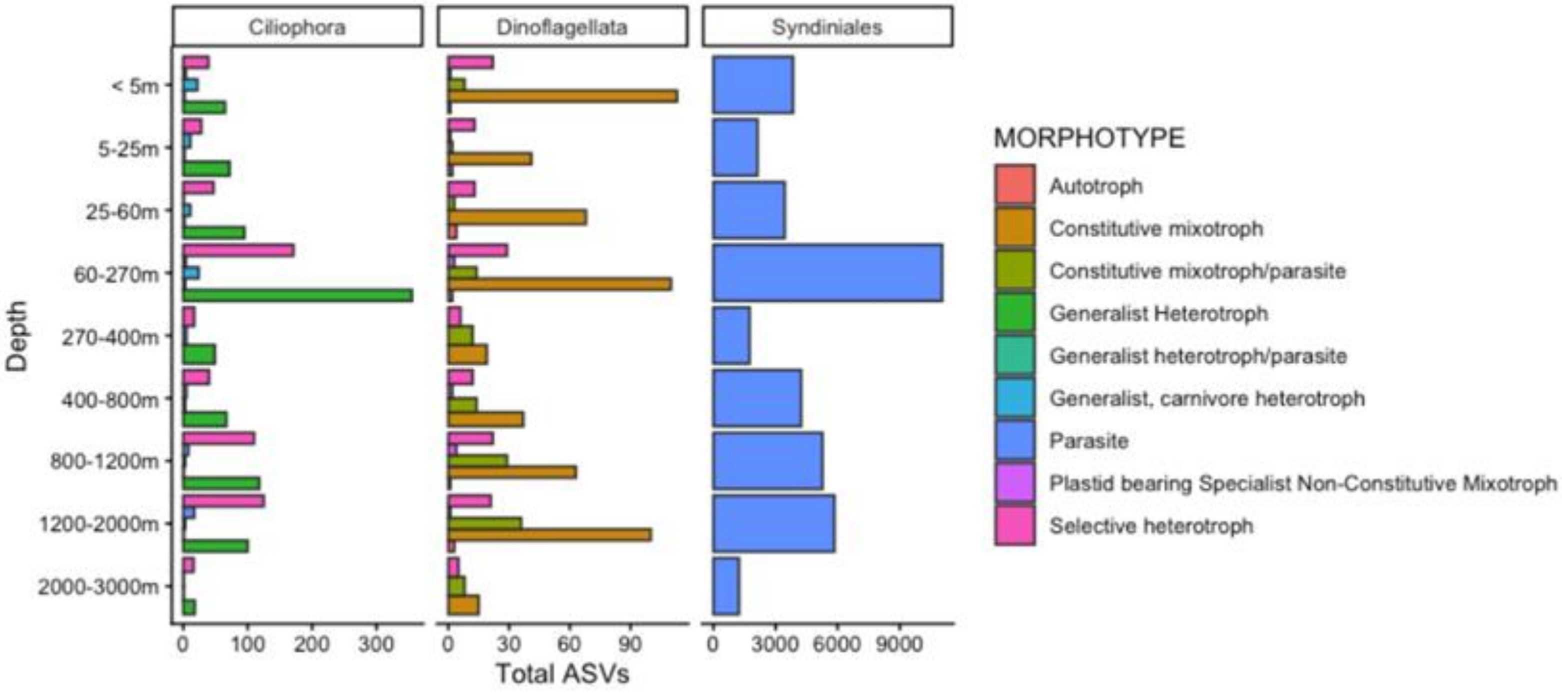


**Figure S16**. Total number of ASVs across ciliates and dinoflagellates (non-syndiniales and syndiniales only) assigned trophic modes (Jones et al. 2025). Bar plots report the specific feeding mode (morphotype) across depth (y-axis).
